# Induction of apoptosis by double-stranded RNA was present in the last common ancestor of cnidarian and bilaterian animals

**DOI:** 10.1101/2023.11.20.567790

**Authors:** Itamar Kozlovski, Adrian Jaimes-Becerra, Ton Sharoni, Magda Lewandowska, Ola Karmi, Yehu Moran

**Author notes:** These authors contributed equally to this work (Itamar Kozlovski, Adrian Jaimes-Becerra). Corresponding authors (IK); (YM).

## Abstract

Apoptosis, a major form of programmed cell death, is an essential component of host defense against invading intracellular pathogens. Viruses encode inhibitors of apoptosis to evade host responses during infection, and to support their own replication and survival. Therefore, hosts and their viruses are entangled in a constant evolutionary arms race to control apoptosis. Until now, apoptosis in the context of the antiviral immune system has been almost exclusively studied in vertebrates. This limited phyletic sampling makes it impossible to determine whether a similar mechanism existed in the last common ancestor of animals. Here, we established assays to probe apoptosis in the sea anemone *Nematostella vectensis*, a model species of Cnidaria, a phylum that diverged approximately 600 million years ago from the rest of animals. We show that polyinosinic:polycytidylic acid (poly I:C), a synthetic long double-stranded RNA mimicking viral RNA and a primary ligand for the vertebrate RLR melanoma differentiation-associated protein 5 (MDA5), is sufficient to induce apoptosis in *N. vectensis*. Furthermore, at the transcriptomic level, apoptosis related genes are significantly enriched upon poly(I:C) exposure in *N. vectensis* as well as bilaterian invertebrates. Our phylogenetic analysis of caspase family genes in *N. vectensis* reveals conservation of all four caspase genes involved in apoptosis in mammals and revealed a cnidarian-specific caspase gene which was strongly upregulated. Altogether, our findings suggest that apoptosis in response to a viral challenge is a functionally conserved mechanism that can be traced back to the last common ancestor of Bilateria and Cnidaria.

## Introduction

Apoptosis, a major form of programmed cell death, is one of the most important innovations of multicellular eukaryotes, and its components, at least in some form, are found in representatives of all multicellular eukaryotes (Koonin and Aravind 2002b). The field of apoptosis research has thrived since the discovery of its molecular mechanism in the nematode *Caenorhabditis elegans* (Yuan, et al. 1993; Yuan and Horvitz 2004). Since then, apoptosis has been extensively studied in mammals, insects, and nematodes, and has also been described in some of the most basally-branching animal lineages, such as sponges and cnidarians (Wiens, Krasko, Blumbach, et al. 2000; Wiens, Krasko, Müller and Müller 2000; Seipp, et al. 2001; David, et al. 2005; Pankow and Bamberger 2007; Oberst, et al. 2008; Lasi, et al. 2010; Pernice, et al. 2011). Furthermore, processes similar to apoptosis have also been observed in plants (Reape and McCabe 2010), fungi (Madeo, et al. 2002; Hamann, et al. 2008; Sharon, et al. 2009), and even in some unicellular eukaryotes (Bidle and Falkowski 2004; Deponte 2008; Pollitt, et al. 2010; Kaczanowski, et al. 2011). In bacteria, other forms of regulated cell death such as pyroptosis have been described (Johnson, et al. 2022) whereas diverse homologs of key components of the apoptotic machinery have been found in bacterial genomes (Koonin and Aravind 2002a; Allocati, et al. 2015). Interestingly, recent studies have demonstrated that bacteria employ regulated forms of cell death in response to phage infection, thereby protecting the nearby population from the spread of infection (Rousset and Sorek 2023). These studies suggest that some of the prokaryotic defense systems involving regulated cell death are evolutionarily ancestral of key components of eukaryotic immune systems and that regulated cell death is an efficient mechanism for blocking viral infection in all domains of life.

In their struggle for existence, viruses and hosts have evolved diverse mechanisms to control and manipulate the apoptotic pathway. Cell death is important for limiting pathogen replication in infected cells, while simultaneously promoting the inflammatory and innate immune responses that are crucial for host immunity (Upton and Chan 2014). Since viruses are obligatory intracellular parasites, they must be able to manipulate the apoptotic pathways in order to control the lifespan of their host and to replicate. Evidently, many viruses encode inhibitors of apoptosis to evade host responses during infection and to support their own replication and survival (Benedict, et al. 2002). Dysregulated cell death as a result of defects in the apoptotic pathways is often a characteristic of inflammatory, autoimmune disorders, and cancer (Bouillet, et al. 1999; van Dommelen, et al. 2006; Carneiro and El-Deiry 2020). Therefore, as both viruses and cells struggle for control of cell death pathways, evolutionary arms race has shaped a complex host-pathogen interaction at this front (Orzalli and Kagan 2017).

Apoptosis has morphological characteristics that include cell shrinkage, nuclear fragmentation, chromatin condensation and membrane blebbing, all of which are the result of the proteolytic activity of the caspase proteases (Li and Yuan 2008; Taylor, et al. 2008). Upon activation, caspases induce apoptosis through the cleavage of several proteins, eventually leading to the phagocytic recognition and engulfment of the dying cell (Tait and Green 2010). In vertebrate cells, apoptosis typically proceeds through one of two signaling pathways termed the intrinsic and extrinsic pathways, both of which result in activation of the executioner caspases, Caspase-3 and Caspase-7 (Tait and Green 2010). The intrinsic pathway is initiated by cell stress that is sensed by internal sensors such as p53, whereas the extrinsic pathway is initiated by an external stimulus such as death ligand/death receptor (for example, TNF and TNF receptor) interactions (Benedict, et al. 2002).

Double-stranded RNA (dsRNA) is associated with most viral infections, either directly by the viral genome itself (in the case of dsRNA viruses) or indirectly by generating intermediates in host cells during viral replication (Son, et al. 2015; Chen and Hur 2022). In mammals, a multitude of studies demonstrated that dsRNA is able to activate a strong antiviral response through the stimulation of type I IFN (IFN-I), and to inhibit the growth of mouse tumors (Barber 2001; Estornes, et al. 2012). In vertebrates, viral RNAs are recognized by intracellular pattern recognition receptors (PRRs) such as the RIG-I-like receptors (RLRs) retinoic acid-inducible gene I (RIG-I) and melanoma differentiation-associated protein 5 (MDA5) (Kato, et al. 2006; Dixit and Kagan 2013; Rehwinkel and Gack 2020). Upon recognition of their ligands, RIG-I and MDA5 interact with the adaptor protein mitochondrial anti-viral signaling protein (MAVS), which provides a scaffold to activate the NF-κB and IRF3 transcription factors (Sun, et al. 2006). Both transcription factors, in turn, regulate the expression of genes that can initiate the apoptotic pathway (Chattopadhyay, et al. 2010b; Orzalli and Kagan 2017).

Until now, apoptosis in the context of the antiviral immune system has been almost exclusively studied in vertebrates. From this limited phyletic sampling, it is impossible to deduce what was the original mode of action of the antiviral response in the last common ancestor of vertebrates and whether apoptosis played a role in antiviral immunity in early-branching animal lineages. The phenomenon of bleaching in Cnidaria (sea anemones, corals, hydroids and jellyfish) has motivated several groups to address questions regarding apoptosis. In the stony coral *Stylophoea pistillata,* it was demonstrated that bleaching and death of the host animal in response to thermal stress involve a caspase-mediated apoptotic cascade induced by reactive oxygen species produced primarily by the algal symbionts (Tchernov, et al. 2011). A study of the stony corals *Pocillopora damicornis* and *Oculina patagonica* demonstrated that reduced pH conditions induce tissue-specific apoptosis that leads to the dissociation of polyps from coenosarcs (Kvitt, et al. 2015). Another study demonstrated that coral cells undergo apoptosis in response to human TNFα and that coral TNF kills human cells through direct interaction with the death receptor pathway (Quistad, et al. 2014). Pyroptosis, a regulated form of inflammatory cell death that involves plasma membrane pore formation mediated by gasdermin family proteins (Bergsbaken, et al. 2009), was recently reported in response to lipopolysaccharide (LPS) in *Hydra vulgaris* (Chen, et al. 2023a) and in response to the bacterial pathogen *Vibrio coralliilyticus* in the reef-building coral species *Pocillopora damicornis* (Jiang, et al. 2020). Nevertheless, the role of apoptosis or other forms of cell death as antiviral mechanisms in invertebrates remains mostly unexplored.

Over the past two decades the sea anemone *N. vectensis* has gained popularity as a model organism for studying molecular evolution, traditionally in the context of development and regeneration (Layden, et al. 2016; Rottinger 2021). As a cnidarian, *N. vectensis* is particularly informative for comparative biology. This is because Cnidaria is a sister group to Bilateria (the group including the vast majority of extant animals) and these two groups diverged approximately 600 million years ago from their last common ancestor (Erwin, et al. 2011; Technau and Steele 2011; Layden, et al. 2016). We have previously shown that *N. vectensis* possesses two RLR paralogs (named RLRa and RLRb) of the mammalian MDA5 that senses dsRNA. Further, we showed that RLRb binds to long dsRNA to initiate a functionally conserved innate immune response (Lewandowska, et al. 2021). In this study, we sought to determine whether and how dsRNA affect apoptosis in *N. vectensis*. We found that the dsRNA viral mimic poly(I:C) is sufficient to strongly induce apoptosis in *N. vectensis* and uncovered a conserved network of genes involved in this process, pointing to a functional conservation of dsRNA-induced apoptosis that existed before the cnidarian-bilaterian split.

## Results

### dsRNA induces apoptosis in *N. vectensis* derived cells *ex-vivo*

We used Apotracker-green (Apo-15), an Annexin V based marker which is commonly used in mammalian cells to probe apoptosis by flow cytometry (Crowley, et al. 2016; Barth, et al. 2020). We first used mitomycin-c (MMC) and cycloheximide (CHX) treatments, common inducers of apoptosis in human and mouse cells (Pirnia, et al. 2002; Baskić, et al. 2006), as positive controls to validate this assay in *N. vectensis*. Using imaging flow cytometry (ImageStream^X^), we observed morphological differences between cells that were negative to Apotracker green and those that were positive to Apotracker green and/or the viability dye Zombie NIR (**Supplementary figure 1**). Treatment of *N. vectensis* derived cells with 2 mM of the translation blocker cycloheximide for 48 hours resulted in a significantly increased Apotracker green fluorescence intensity compared to DMSO (p=0.001269, n=3, two-sided t-test) (**Supplementary figure 1B, 2A, 2B**). Notably, cells that were negative to Apotracker green had a significantly higher circularity score compared to Apotracker positive cells, and compared to cells that were positive to both Apotracker and Zombie NIR, the later had the lowest score (**Supplementary figure 1C**). MMC, an antitumor drug known to induce apoptosis by crosslinking DNA (Tomasz 1995; Pirnia, et al. 2002), was previously used to kill proliferating cells before stem cell transplantation in *N. vectensis* (Talice, et al. 2023). Indeed, zygotes treated with 60 µM MMC for 48 hours had a significantly increased cell death compared to their DMSO counterparts (p=0.002472, n=4, two-sided t-test) (**Supplementary figure 2C, 2D**). Next, we studied the effect of poly(I:C) on apoptosis. In mammals, poly(I:C) is recognized, among other intracellular receptors, by the extracellular toll-like receptor 3 (TLR3) which is expressed on the cell surface of fibroblasts and epithelial cells (Matsumoto and Seya 2008). *N. vectensis* has only one toll-like receptor (TLR), which is not orthologous of TLR3. This receptor was shown to respond to bacterial pathogens and its exact ligand is still unknown, however, was suggested to be Flagellin (Brennan, et al. 2017). Indeed, injection of poly(I:C) into *N. vectensis* zygotes resulted in strong upregulation of RLRa and RLRb (**Supplementary figure 2E**), whereas incubation of zygotes with high concentrations of poly(I:C) (0.5 µg/µL) did not increase the expression of neither RLRa nor RLRb (**Supplementary figure 2F**), suggesting that *N. vectensis* can only respond to intracellular dsRNA. To deliver poly(I:C) into cells, we transfected *N. vectensis*-derived cells with 2 µg/ml poly(I:C). Indeed, we detected a significantly increased Apotracker green fluorescence intensity in poly(I:C) transfected cells relative to their mock transfected counterparts (p=0.0265, n=3, one-sided t-test) (**Figure 1A, 1B**).

**Figure 1.**
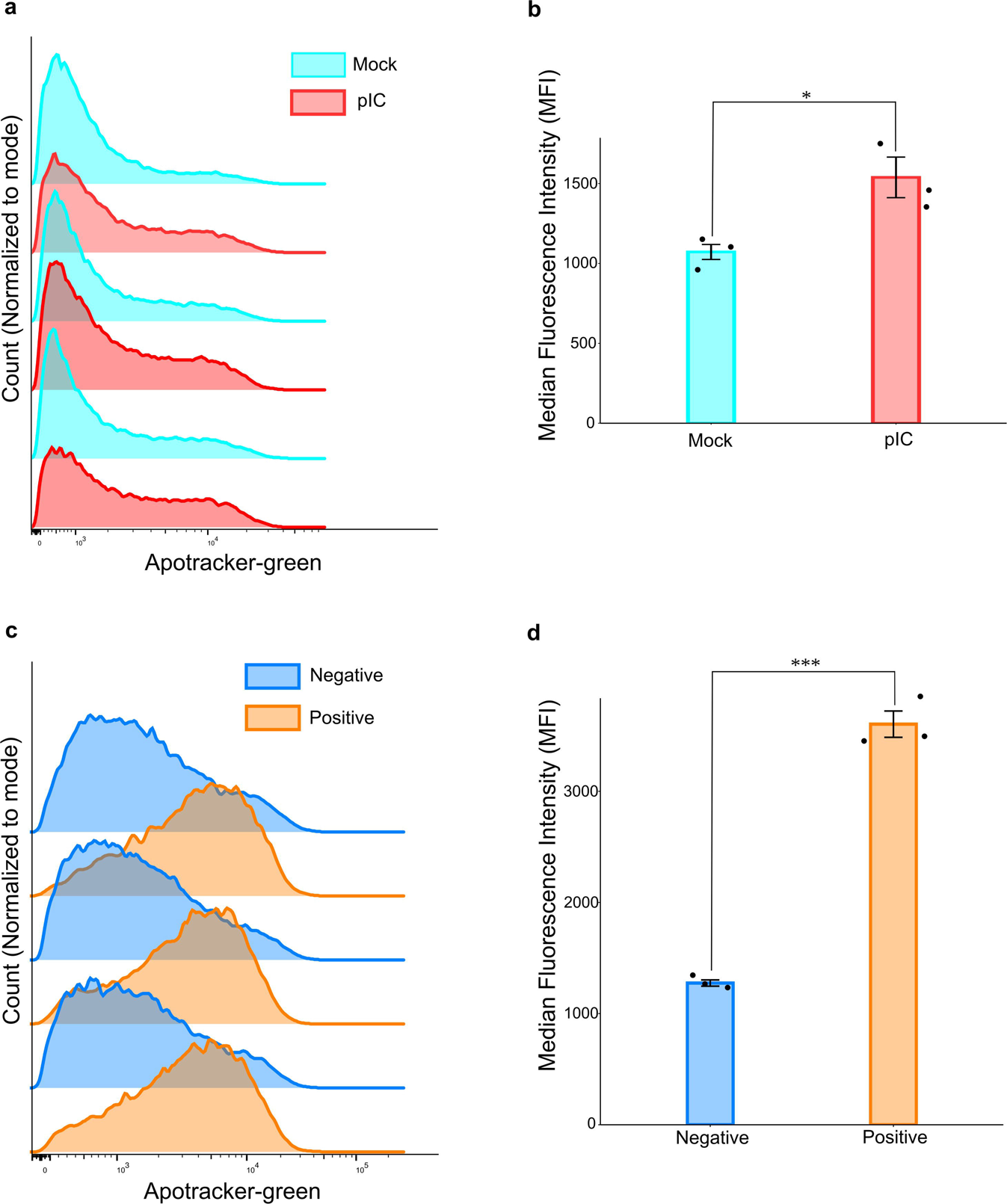
poly(I:C) induces cell death *ex vivo*. (a) Apotracker green was used to quantify apoptosis 24 hours following 2 ug/ml poly(I:C) transfection in *N. vectensis* derived cells. Fluorescence intensity determined by flow cytometry is shown on a logarithmic scale in arbitrary units on the X axis. The histograms were normalized to the mode. Mock transfected cells are shown in light blue and poly(I:C) transfected cells are shown in red. (b) Quantification of the data shown in (a). The bars represent median fluorescence intensity (MFI). Error bars represent standard deviation. (c) *N. vectensis* derived cells were transfected with poly(I:C) labeled with rhodamine. Rhodamine negative cells (shown in blue) were then compared to rhodamine positive cells (shown in orange). (d) Quantification of the data shown in (c). The bars represent median fluorescence intensity (MFI). Error bars represent standard deviation. Three biological replicates were used for each condition. 30,000 events were recorded per sample. All comparisons were done by one sided t-test. Individual data points are shown as a jitter. * p<0.05, **p<0.01, ***p<0.001.

As of today, no cytokines have been described in *N. vectensis*. Thus, it is unclear whether infected cells are capable of alarming neighbors upon viral infection (i.e., paracrine signaling), or alternatively, the signaling is limited to the infected cells (i.e., autocrine signaling). To address this question, we transfected cells with poly(I:C) labeled with rhodamine. Flow cytometry revealed transfection efficiency ranging from about 29 % to 44 % (**Supplementary figure 3A-C**). Next, we asked whether cells that were successfully transfected with poly(I:C) (rhodamine positive) were prone to apoptosis more than cells that did not uptake the labeled poly(I:C). Indeed, the Apotracker-green signal was increased in cells that were also positive to rhodamine (p=0.00081, n=3, one-sided t-test) (**Figure 1C, 1D**), indicating that poly(I:C) triggered the apoptosis mostly in cells that internalized it.

### dsRNA induces apoptosis in *N. vectensis* injected animals

Caspases are key mediators of apoptosis. Among them, Caspase-3 is a major effector caspase involved in the execution phase of apoptosis (Nicholson, et al. 1995). Following cleavage and activation by initiator caspases (CASP8, CASP9 and/or CASP10), the mammalian Caspase-3 executes apoptosis by catalyzing cleavage of many essential key proteins (Nicholson, et al. 1995; Walsh, et al. 2008; Nakatsumi and Yonehara 2010; Thomsen, et al. 2013). Methods for determining Caspase-3 activity are based on the proteolytic cleavage of its preferred recognition sequence Asp-Glu-Val-Asp (DEVD) (Gurtu, et al. 1997). Such methods have been applied for studying apoptosis upon stress conditions in other cnidarians including corals and the sea anemone *Exaiptasia diaphana* (Pernice, et al. 2011; Bieri, et al. 2016; Ros, et al. 2016). We compared the Caspase-3 activity using a colorimetric assay in poly(I:C) injected animals versus NaCl control at different time points. At 6 hours post injection (hpi), we did not observe a significant difference in Caspase-3 activity compared to NaCl control (p=0.56, n=3, one-sided t-test) (**Figure 2A**), whereas caspase activity was significantly increased at 24 hpi (p=0.0013,n=3, one-sided t-test) and did not reach a statistical significance at 48 hpi (p=0.069,n=3, one-sided t-test) (**Figure 2B, 2C**). The increase in Caspase-3 activity relative to 6 hours, although did not reach statistical significance, was most noticeable at 24 hours (p=0.145, n=3, one way ANOVA test) after injection and decreased in magnitude at 48 hpi (p=0.469, n=3, one way ANOVA test) (**Figure 2D**).

**Figure 2.**
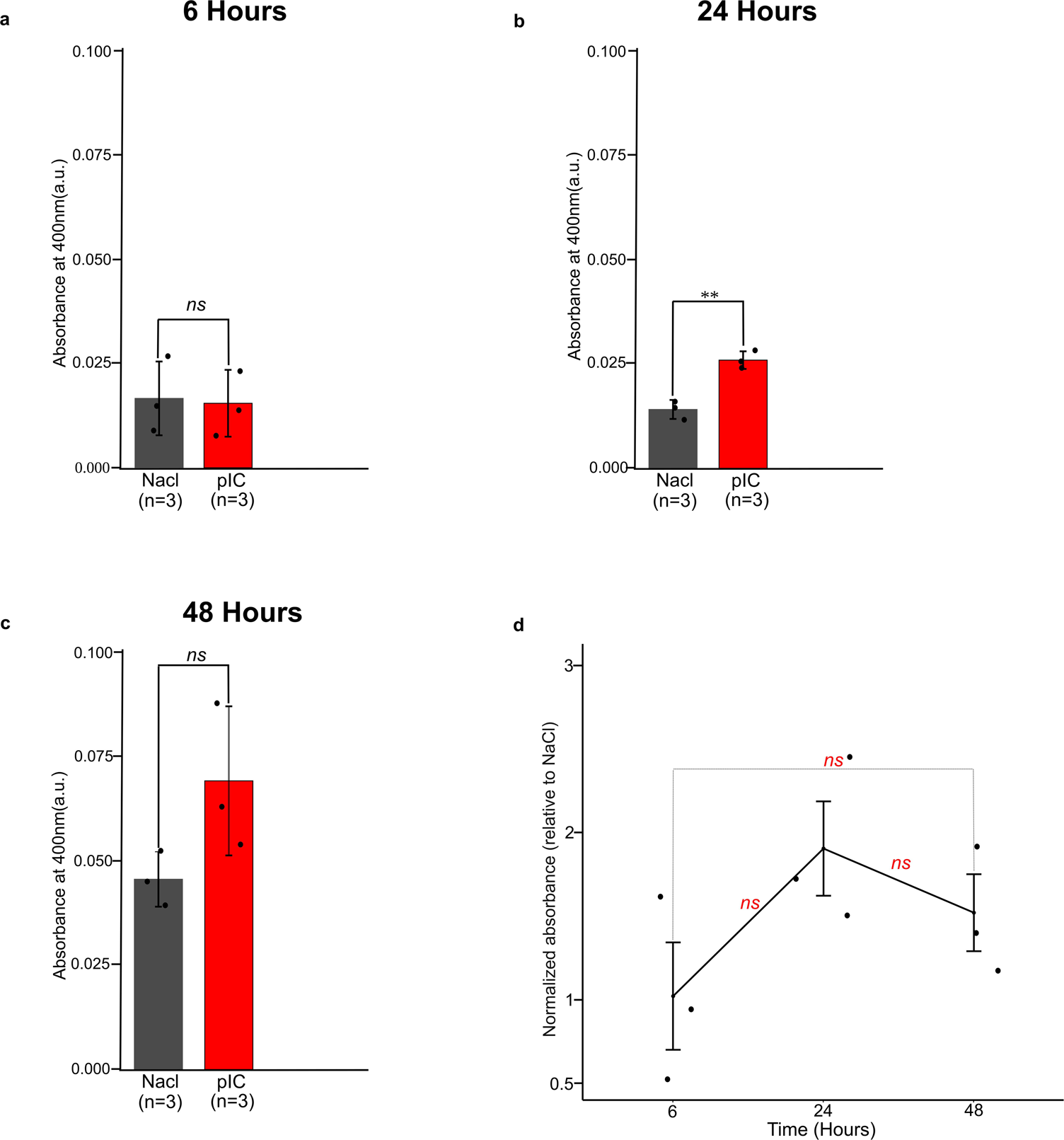
poly(I:C) increases caspase-3 activity *in-vivo*. Zygotes were injected with poly(I:C) and subjected to caspase-3 activity colorimetric assay. Optical density was measured at 400nm. (a) Lysates were obtained from either poly(I:C) or NaCl control treated embryos at 6 hours post injection (hpi), (b) at 24 hours post injection, and (c) at 48 hours post injection. (d) Optical density in poly(I:C) treated animals (relative to its NaCl treated controls) at different time points. Three biological replicates were used for each condition. For panels (a)-(c) one-sided t test was performed. For panel (d) one way ANOVA test with Tukey’s post hoc test was performed. Individual data points are shown as a jitter. * p<0.05, **p<0.01, ***p<0.001.

### dsRNA triggers the activation of genes associated with the apoptotic pathway

To further investigate the impact of dsRNA on apoptosis, we reanalyzed a gene expression dataset previously published by Lewandowska et al. (2021), which examined the antiviral immune response of *N. vectensis* to a viral dsRNA mimic (poly(I:C)). Compared with 0.9% NaCl control, dsRNA induced a gene expression pattern strongly associated with apoptotic process including upregulated expression of genes encoding initiator and executioner caspases (Caspase-8 and Caspase-3/6/7), negative apoptotic regulators (Bcl, MCL and IAP), and positive apoptotic regulators (Bok, Apaf-1 and TRAFs) for 24 hpi. Consistent with the caspase-3 activity assay **(Figure 2D)**, this increase in gene expression was transient as a reduction in the number of upregulated apoptosis-related genes was observed within 48 hpi. No upregulated apoptosis-related genes were detected at 6 hpi (**Figure 3A, Supplementary file 1**). Gene ontology (GO) term analysis revealed significant enrichment of apoptosis-related terms induced by dsRNA (**Figure 3B**), with the term ‘Regulation of apoptotic process’ being most significant at the 24 hpi time point. Furthermore, based on the identified upregulated genes associated with apoptosis in *N. vectensis* at 24 hpi, it seems that both the intrinsic and the extrinsic pathways of apoptosis were activated. The intrinsic pathway (depicted on the left in **Figure 3C**) may be initiated by various intracellular signals, such as DNA damage. These signals lead to the downregulation of pro-survival proteins like Bcl-2 and Bcl-10, thereby enabling pro-apoptotic molecules such as Bok, to provoke a mitochondrial permeability transition. This transition results in the release of cytochrome c and other pro-apoptotic molecules, counteracting the anti-apoptotic activity of IAPs. Following this, cytochrome c associates with Apaf-1 to form the apoptosome, which activates initiator caspases. Subsequently, initiator caspases trigger the executioner caspase cascade, culminating in cell death. The extrinsic pathway (**Figure 3C**) is initiated by the binding of secreted ligands from the TNF family to their corresponding receptors within the TNFR family. Upon activation, receptor homotrimers or homomers recruit cytoplasmic adaptor molecules, which then instigate cell death through various pathways. Following the activation of classical cell death receptors (e.g., FAS, DR4/5, TNFR1), adaptor molecules directly bind and facilitate the cleavage of initiator caspases. These, in turn, activate the effector caspases that orchestrate cell death (Bertheloot, et al. 2021).

**Figure 3.**
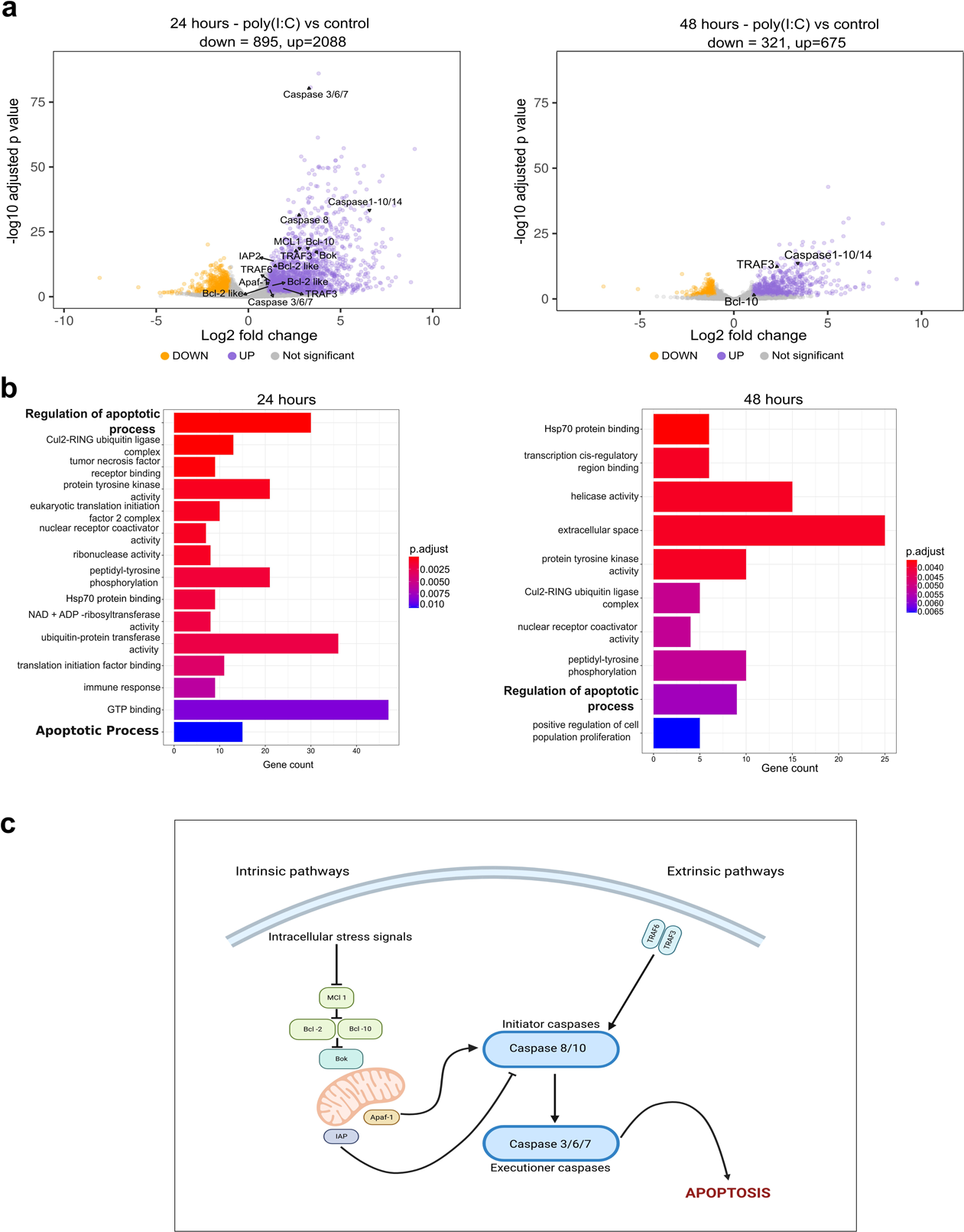
Micronjection of the dsRNA viral mimic poly(I:C) induces activation of apoptosis pathway genes. (A) Volcano plots showing differentially expressed genes (DEGs) in *N. vectensis* after poly(I:C) administration at 24 hpi and 48 hpi. Significantly up- and downregulated genes (adjusted P-value < 0.05 and absolute log_2_ fold change > 1) are highlighted in purple and orange, respectively. Among those, genes associated with the apoptosis pathway are highlighted, along with their names. (B) The bar plots illustrate enriched Gene Ontology (GO) terms, showcasing the Top 15 and Top 10 overrepresented biological process GO terms among up-regulated genes at the 24 hpi and 48 hpi time points, respectively. All terms displayed are statistically significant as per the hypergeometric test (with a Benjamini-Hochberg corrected P-value ≤ 0.05). (C) A simplified schematic illustrates the proposed intrinsic and extrinsic apoptotic signaling pathways in *N. vectensis* based on the components depicted are up-regulated post poly(I:C) administration at 24 hpi and possess homologs in their metazoan counterparts.

### Caspases in Metazoa: Phylogenetic analysis and domain organization

Six putative caspase genes were identified in the genome of *N. vectensis*. A phylogenetic analysis was conducted to explore the distribution of these caspase genes from *N. vectensis* across Metazoa. The Maximum-Likelihood tree depicted in **Figure 4** shows that the caspases from *N. vectensis* are grouped into five distinct clades, with the exception of clade II (highlighted in green), where Caspase-9 is exclusively found in deuterostome animals. All caspases within this clade possess a CARD and CASc domain; in humans, they are recognized as initiator caspases (Li, et al. 1997). The sequences in the tree were named based on their human orthologs. Clade I exclusively comprises sequences from Anthozoa, and showcases a domain organization with both NHR and CASc domains. The Neuralized Homology Repeat (NHR) domain, a module of approximately 160 amino acids, was initially identified as a tandem repeat in the *Drosophila* Neuralized, a protein instrumental in the development of the central and peripheral nervous systems (Nakamura, et al. 1998). Intriguingly, within this clade, we discovered a sequence from *N. vectensis* (NVE5282) that is the most upregulated at 24 hpi time point, exhibiting a Log2 Fold Change (LFC) of 6.4. Clade III, highlighted in red, encompasses sequences from Protostomia, Deuterostomia, Medusozoa, and Anthozoa lineages. A defining feature of the caspases in this clade is the presence of dual death-effector domains (DEDs), which serve as recruitment domains, alongside CASc domains. Caspases with this domain structure (Caspase-8 and −10 in humans) are identified as initiator caspases in the apoptotic pathway (Lamkanfi, et al. 2002). Within this clade, a sequence from *N. vectensis* (NVE9681) was found to be upregulated at the 24 hpi time point, with a LFC of 2.8. Clade IV, highlighted in yellow, displays the greatest diversity regarding the number of human orthologs present as well as domain structure. This clade includes sequences from the Protostomia, Deuterostomia, and Anthozoa lineages. Within this clade, we observe subclades wherein inflammatory caspases from humans (Caspase-1, −4 and −5) are located. Adjacent to this subclade, human Caspase-2, identified as an initiator caspase, along with Caspase-14, recognized for its functional role in development, are also located (Sakamaki and Satou 2009). In this clade, there is a sister group to the remaining species, exclusively consisting of anthozoan sequences, which possess both CARD and CASc domains.. Within this group, a sequence from *N. vectensis* (NVE26090) is found, however, it is not upregulated at any time point in our experiment. This clade exhibits low bootstrap support, yet high SH-aLRT and aBayes support. This discrepancy could arise from a matter of statistical sensitivity, wherein SH-aLRT and aBayes tests may be more sensitive to certain data patterns. They might detect signals in the data supporting the branch that bootstrap analysis might miss, especially if the dataset is small or the signal is weak (Guindon, et al. 2010). Clade V stands as the only group devoid of human orthologs, predominantly featuring sequences from Anthozoa and Medusozoa. The protein domain structure within this clade is characterized by the sole presence of the CASc domain. Within this clade, two sequences from *Hydra vulgaris* appear in bold. Recently, these sequences were discovered to have an inflammatory function (Chen, et al. 2023b). Within this clade, a sequence from *N. vectensis* (NVE20429) is found, which is not upregulated at any time point in our experiment. Clade VI represents the largest clade both in terms of the number of sequences and the diversity of lineages. It encompasses sequences from Porifera, Placozoa, Protostomia, Deuterostomia, Medusozoa, and Anthozoa. In this clade, we identified a subclade whose sequences are characterized solely by the presence of the CASc domain, and exclusively include sequences from the Protostomia and Deuterostomia lineages. The human Caspase-3, −6, and −7, grouped within this subclade, are identified as executioner caspases in the apoptotic pathway (Sakamaki and Satou 2009). The remaining subclades, apart from possessing the CASc domain, also feature an N-terminal CARD domain. Within these subclades, we identified two upregulated sequences from *N. vectensis*: NVE23160 with a LFC of 1.15 and NVE21851 with a LFC of 3.37 at 24 hpi. It is noteworthy that the tree we obtained using Bayesian inference is similar to this obtained through the maximum likelihood method (**Supplementary Figure 4**).

**Figure 4.**
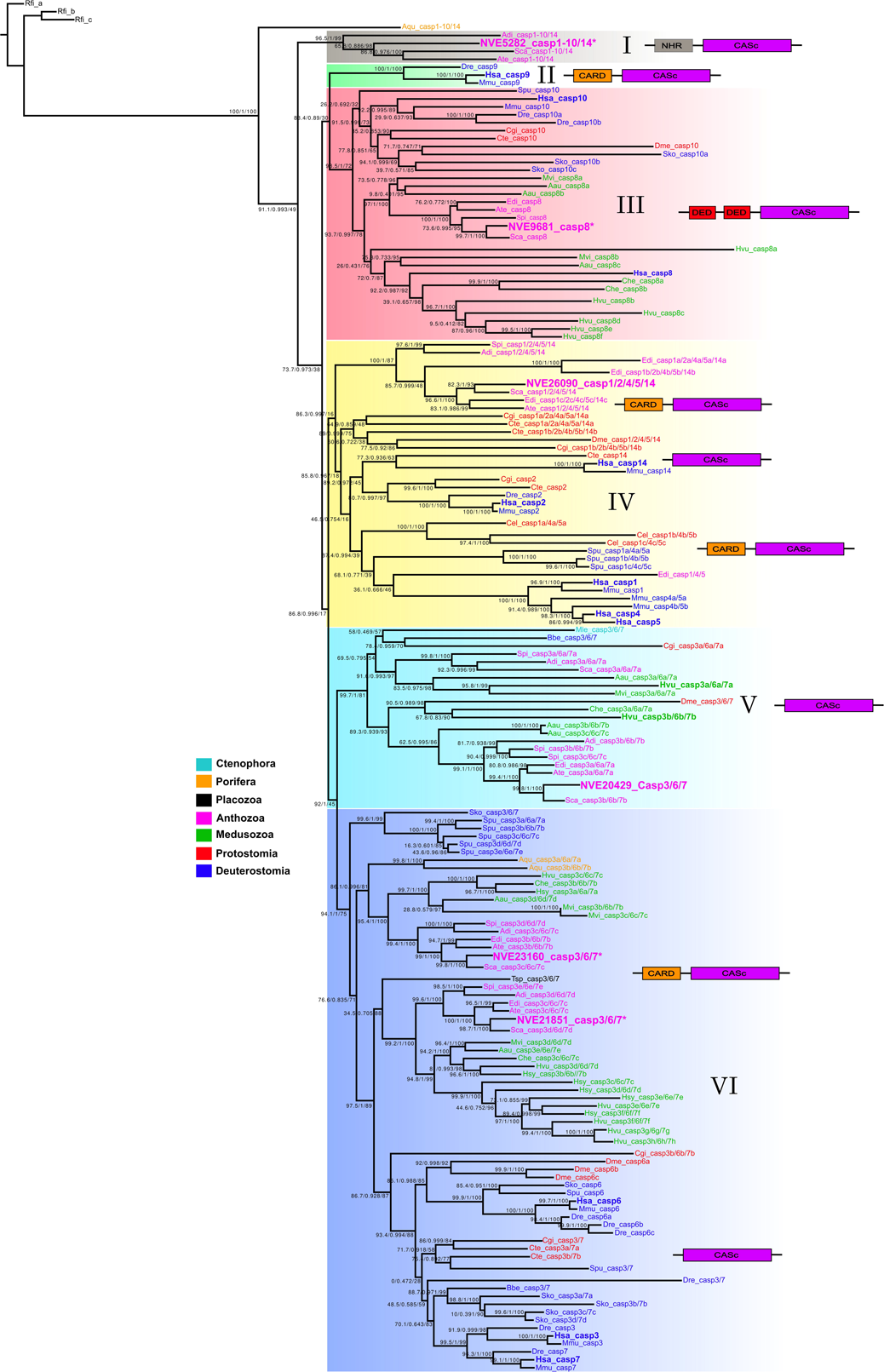
Phylogenetic relationship among metazoan caspases. Numbers on nodes, separated by slashes, represent SH-aLRT, approximate Bayes test, and ultrafast bootstrap support values, in that order, for Maximum Likelihood (ML) analysis. The domain organization is displayed alongside each clade. CASc: caspase domain, CARD: caspase-recruitment domain, DED: death-effector domain, NHR: neuralized homology repeat. Sequences from *N. vectensis* are highlighted in bold magenta and presented in a larger font. Some of these sequences feature an asterisk, indicating that these genes are upregulated at the 24hpi time point with a Log2 Fold Change > 1. *Homo sapiens* sequences are highlighted in bold blue. Rfi: *Reticulomyxa filosa*, Aqu: *Amphimedon queenslandica*, Adi: *Acropora digitifera*, NVE: *N. vectensis vectensis*, Sca: *Scolanthus callimorphis*, Ate: *Actinia tenebrosa*, Dme: *Drosophila melanogaster*, Dre: *Danio rerio* Hsa: *Homo sapiens*, Mmu: *Mus musculus*, Spu: *Strongylocentrotus purpuratus*, Cgi: *Crassostrea gigas*, Cte: *Capitella teleta*, Sko: *Saccoglossus kowalevskii*, Mvi: *Morbakka virulenta*, Aau: *Aurelia aurita*, Edi: *Exaiptasia diaphana*, Spi: *Stylophora pistillata*, Hvu: *Hydra vulgaris*, Che: *Clytia hemisphaerica*, Cel: *Caenorhabditis elegans*, Mle: *Mnemiopsis leidyi*, Bbe: *Branchiostoma belcheri*, Tsp: *Trichoplax sp.*, Hsy: *Hydractinia symbiolongicarpus*.

Recently, a study that encompasses a comprehensive phylogenetic analysis of the caspase family was published (Krasovec, et al. 2023). This study presents certain parallels with our study, especially in the identification of specific monophyletic groups. For instance, both analyses recognize a group, which we term Group III, characterized by its dual Death Effector Domains (DED). Nonetheless, there are significant differences as well. While the Krasovec et al. study recovers caspase groups based on domain structure, our topology diverges from this pattern. This discrepancy likely stems from differing taxonomic sampling strategies. Our analysis, in contrast to Krasovec et al. which uses 76 sequences, incorporates 163 sequences from various major metazoan lineages, with a particular emphasis on the Cnidaria phylum. These methodological differences highlight the distinct perspectives each study brings to understanding the evolution of the caspase family. Additionally, our phylogenetic analysis primarily aims to ascertain the placement of different *N. vectensis* genes within this evolutionary context.

### dsRNA induces a conserved antiviral response pathway across metazoans

To obtain a wider picture on the evolution of the antiviral apoptotic response in animals, we analyzed data from seven studies (Zhang, et al. 2017; Hagai, et al. 2018; Linehan, et al. 2018; Lafont, et al. 2020; Lewandowska, et al. 2021; Bar Yaacov 2022; Tassia, et al. 2023) focused on the gene expression response to immune challenges. These studies specifically employed poly (I:C) as a viral mimic. Importantly, Bar Yaacov’s 2022 study on the planarian *Schmidtea mediterranea* employed double-stranded RNA virus, rather than poly (I:C), as the immunological challenge. Despite differences in the species studied, experimental designs, and analytical methods, these studies collectively contribute to our understanding of dsRNA’s influence on apoptosis on an evolutionary scale. Based on the results of the gene ontology (GO) analysis conducted in each study, we found that GO terms related to apoptosis are significantly enriched in all the studies, except for the hemichordate *Saccoglossus kowalevskii*, which did not show any significant GO terms related to apoptosis (**Figure 5**). Similarly, under the apoptosis categories, we pinpointed clusters of upregulated genes within crucial protein families in the apoptotic pathway, including caspases (both initiators and executioners), anti-apoptotic regulators like the Bcl-2 family members, and pro-apoptotic factors such as Apoptotic Protease-Activating Factor 1 (Apaf-1), Tumor Necrosis Factor (TNFs), and TNF Receptor Associated Factor (TRAFs) (**Figure 5**). The majority of lineages exhibited a shared upregulation of these genes.

**Figure 5.**
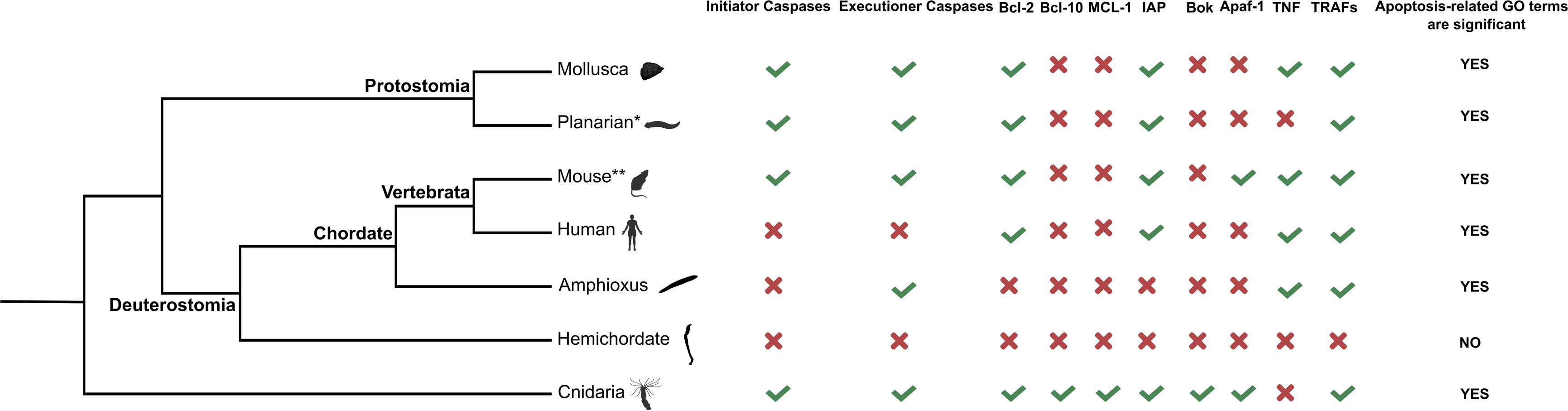
A simplified phylogenetic tree representing selected groups, highlighting the regulatory patterns of key apoptosis gene families. In this representation, a check mark (√) denotes the up-regulation of a gene, while an X indicates the absence of upregulation following exposure to the dsRNA viral mimic poly(I:C) or virus. *The study pertaining to Platyhelminthes utilized a dsRNA virus as opposed to poly(I:C). **This information is synthesized from two separate studies. The specific details of each study are available in supplementary file 3.

## Discussion

The co-evolution of viruses with their hosts has led to the emergence of viral pathogens that are capable of evading or actively suppressing host immunity (Chan and Gack 2016). The ability to recognize and respond to foreign nucleic acids among an abundance of self nucleic acids in an organism is, therefore, crucial for host fitness (Schlee and Hartmann 2016). Long dsRNA in the cytosol is a hallmark of DNA and RNA virus replication and is absent from an uninfected host cell (Weber, et al. 2006; Son, et al. 2015). In vertebrates, poly(I:C), which mimics dsRNA, binds to and triggers the activation of the RNA sensors PKR, OAS1, TLR3, MDA5 and RIG-I, and consequently poly(I:C) or its derivatives have been used as a tool to identify and characterize dsRNA recognition receptors and ligand requirements (Alexopoulou, et al. 2001; Yoneyama, et al. 2004; Gitlin, et al. 2006; Kato, et al. 2006; Kato, et al. 2008; Goubau, et al. 2014). Following recognition of poly(I:C), MDA5 signals via MAVS and IRF3-IRF7 to induce type I INFs which enables infected cells to “alert” neighboring cells against incoming infection and recruit cells of the immune system to battle the virus (Stetson and Medzhitov 2006; Ivashkiv and Donlin 2014). Additionally, MAVS activation promotes apoptosis through several pathways (Besch, et al. 2009; Chattopadhyay, et al. 2010a; Glas, et al. 2013; El Maadidi, et al. 2014; Kumar, et al. 2015).

In contrast to vertebrates, most invertebrates, such as insects, plants and nematodes, lack interferons and are thought to rely mainly on RNA interference (RNAi) for antiviral defense (Schlee and Hartmann 2016). However, limited phyletic sampling makes it impossible to determine what was the original mode of action of these systems in their last common ancestor (Iwama and Moran 2023) and whether apoptosis was also induced by viral challenges in early animals. We have previously shown that *N. vectensis* responds to poly(I:C) by upregulating immune-related genes (Lewandowska, et al. 2021) but it remained unclear whether the observed gene expression changes are associated with a phenotype. Here, we filled that gap by showing that *N. vectensis* is capable of inducing apoptosis by exposure to dsRNA alone, both *ex-vivo* (**Figure 1**) and *in-vivo* (**Figure 2**). In fact, regulation of apoptosis was the most significantly enriched set of genes in GO analysis (**Figure 3**). Our phylogenetic analysis (**Figure 4**) identified orthologs for all four caspase genes responsible for apoptosis in mammals, out of which 3 were upregulated upon poly(I:C) treatment.

Our findings challenge the dichotomy found in the literature between the antiviral immune systems of vertebrates and invertebrates as it seems that *N. vectensis* is capable of activating apoptosis as a response to dsRNA challenge. Our transcriptomic analysis (**Figure 3**) indicated that both intrinsic and extrinsic pathways are activated in response to dsRNA. This result is in agreement with our cross species comparative transcriptomic analysis **(Figure 5**) that demonstrated that (1) most representative metazoans (6 out 7 analyzed) respond to dsRNA by significantly upregulating genes related to apoptosis, and (2) the upregulated genes were mostly conserved between distant species. However, in some lineages it seems that the apoptotic response to dsRNA was lost. For instance, the hemichordate *Saccoglossus kowalevskii* did not exhibit a transcriptomic response to poly(I:C). Similarly, in the nematode C. *elegans*, which has been extensively studied in the context of apoptosis, dsRNA injection did not induce apoptosis (Adamo, et al. 2012). It is possible that apoptosis was decoupled from dsRNA in C. *elegans* as a consequence of a highly efficient RNAi response.

In recent years, a growing body of evidence indicate that other forms of cell death are also involved in the antiviral response. In particular, necroptosis, a regulated form of necrotic cell death (Pasparakis and Vandenabeele 2015), and pyroptosis, a regulated form of inflammatory lytic cell death mediated by gasdermin proteins (Verdonck, et al. 2022), are important host defense strategies for eliminating viral infected cells. The later was observed in cnidaria in the context of antibacterial response (Jiang, et al. 2020; Chen, et al. 2023a). Our phylogenetic analysis revealed that the *N. vectensis* gene (NVE20429) orthologous to the *Hydra vulgaris* gene (**Figure 4**) which was recently reported to be involved in pyroptosis was not upregulated upon poly(I:C) treatment. However, a Gasdermin-encoding gene (NVE20414) was upregulated (LFC=1.8, p=0.0014) at 24 hpi. Importantly, gasdermins are regulated post-translationally by proteolytic cleavage (Xia, et al. 2020). Therefore, based on our data we cannot exclude the possibility that pyroptosis is also activated in response to dsRNA. Further research is required to determine whether pyroptosis and apoptosis are activated simultaneously or whether these two processes are mutually exclusive in *N. vectensis*.

In conclusion, our findings in *N*. *vectensis*, together with our cross-species analysis in the current study, suggest that induction of apoptosis by dsRNA is an ancient response that was present already in the last common ancestor of cnidarian and bilaterian animals, that lived 600 million years ago.

## Methods

### Sea anemone culture

*N. vectensis* polyps were grown in 16 ‰ sea salt water (Red Sea, Israel) at 18 °C. Polyps were fed with *Artemia salina* nauplii three times a week. Spawning of gametes and fertilization were performed according to a published protocol (Genikhovich and Technau 2009) as follows: in brief, temperature was raised to 25 °C for 9 h and the animals were exposed to strong white light. Three hours after the induction, oocytes were mixed with sperm to allow fertilization.

### Cell dissociation

Animals were dissociated into single cells as previously described (Admoni, et al. 2020). Briefly, planulae were washed twice with calcium/magnesium free and EDTA free artificial seawater (17 mM Tris-HCl, 165 mM NaCl, 3.3 mM KCl and 9 mM of NaHCO_3_; final solution pH 8.0) and incubated with 50 µg/ml liberaseTM (Roche, Switzerland) at room temperature for 5-10 minutes with occasional gentle pipetting, until fully dissociated. The reaction was stopped by adding 1/10 volume of 0.5 M EDTA solution. The suspension was filtered using a 35 µm cell strainer. Cells were then centrifuged at 500 × *g* at 4 °C and resuspended in 1× calcium/magnesium free sterile PBS (Hylab, Israel). Cells were counted on hemocytometer and viability was determined using trypan blue (Thermo Fisher Scientific, USA).

### Primary cell culture

Animals were grown at 25 °C for 48 hours post fertilization and dissociated into single cells as described in the previous section. Cells were washed 3 times in PBS before seeding. Viability was assessed using trypan blue. Cell suspension with viability of more than 80 % was used for cell culture. 200,000 cells were seeded per well in a 24 well plate. Cells were grown at 25 °C in L-15 medium (Biological Industries, Israel) supplemented with 10 % FBS, 1 % Pen-Strep, and 2 % 1M HEPES buffer solution.

### Transfection

Transfection was done using lipofectamine 3000 (Thermo Fisher Scientific) according to the manufacturer’s instructions, using high dose lipofectamine.

### Flow cytometry

FACSAria III (BD Biosciences, USA) equipped with 405 nm, 408 nm, 561 nm and 633 nm lasers was used to quantitively assess apoptosis. Per run, 30,000 events were recorded. Helix NP (Ex430/Em470) (Biologend, USA) was used to determine viability. To monitor apoptosis, Apotracker-green (Ex488/Em520) (BioLegend) was used according to the manufacturer’s instructions. FCS files were further analyzed using FlowJo V10 (BD Biosciences, USA). Each measurement consisted of 3 biological replicates, unless indicated otherwise.

### Imaging cytometry

Dissociated cells treated with either CHX or DMSO control were assessed quantitatively using an Amnis® ImageStream^X^ Mk II apparatus (Luminex, USA) equipped with 405nm, 488nm, 642nm and 785nm (SSC) lasers and 6 acquisition channels, using a 60X magnification objective, with low flow rate/ high sensitivity using INSPIRE200 software (Luminex, USA). The INSPIRE200 software was set up using the following parameters: Channel 01 (bright field), Channel 02 for detecting laser 488nm, and Channel 05 detecting laser 642nm. Dissociated cells were stained with 200nM Apotracker green (Ex488/Em520) (Biolegend) and with 1:100 dilution of the Zombie NIR (Ex642/Em746) fixable viability dye (Biolegend) for 15 minutes at room temperature. Cells were then washed twice with PBS and resuspended in 100uL of PBS in 1.5 ml Eppendorf tubes. About 10^7^ cells per sample were used for data acquisition. A total of 10,000 single events were acquired within the focused singlet gate. Focused events were determined by the Gradient RMS parameter for Ch01 and single cells were determined by plotting the area of the events (X-Axis) vs. aspect ratio (Y-Axis) for Ch01. All subsequent analysis was done on this population of cells. Image analysis was run using IDEAS® 6.3 software (Luminex, USA), circularity feature was determined using the Shape Change Wizard, which provided circularity score for each population by measuring the degree of the mask’s deviation from a circle.

### Caspase3 activity assay

For each condition (NaCl vs. poly(I:C)), about 150 injected animals were snap frozen in liquid nitrogen and stored in −80 °C. Caspase-3 Assay Kit (colorimetric) (abcam, UK) was used according to the manufacturer’s instructions. Briefly, cell pellet was lysed in lysis buffer using a homogenizer. Protein quantification was done using BCA assay (Cyanagen, Italy). Same amounts of protein were loaded into each reaction, ranging from 30 µg to 40 µg depending on the yield. Absorbance was measured at 400 nm after 2 hours using a microplate reader (BioTek, USA).

### Poly(I:C) microinjection

To stimulate the antiviral immune response in *N. vectensis*, we used poly(I:C) (Invivogen, USA) as dsRNA viral mimic. We used 3.125 ng/ml of high molecular weight (HMW) poly(I:C) in 0.9 % NaCl (with an average size of 1.5–8 kb), and 0.9 % NaCl as a control. This concentration was determined as sub-lethal in previous titration experiments (Lewandowska, et al. 2021). In each experiment 100–150 zygotes were microinjected and flash frozen in liquid nitrogen at 6 hours, 24 hours, and 48 hours after injection, and then stored at - 80 °C until processed.

### Mitomycin-c and cycloheximide treatments

Zygotes were de-jellified using 3 % cysteine (Merck Millipore, USA), washed 6 times in *N. vectensis* medium (16 ‰ artificial sea water), and incubated with either 60 µM mitomycin-c (Sigma-Aldrich, USA), 2 mM cycloheximide (Abcam, United Kingdom), or an equivalent concentration of DMSO in 24-well plate containing 500 µL *N. vectensis* medium. 200-500 zygotes were grown per well. 48 hours later cells were dissociated from planulae as described in previous sections, stained with Apotracker-green and subjected to flow cytometry. For *ex vivo* experiments, cells were dissociated and seeded in 24-well plates as described in *Primary cell culture* section. 2 mM cycloheximide (Abcam, Cambridge, United Kingdom) or an equivalent concentration of DMSO (Sigma-Aldrich) was used to treat dissociated cells for 48 hours. Cells were then subjected to flow cytometry as described in previous sections.

### RT-qPCR

To test the effect of extracellular poly(I:C), 50 zygotes were incubated with 0.5 µg/µL of poly(I:C) in a total volume of 1 ml 16 ‰ sea salt water (growth medium) (Red Sea, Israel) for 24 hours at 24 °C. as controls, 50 zygotes were incubated only with growth medium. The animals were harvested and snap frozen in liquid nitrogen 24 hours post fertilization. We compared this result to poly(I:C) injection as follows: 200 zygotes were injected with 3.125 ng/µL of poly(I:C) and snap frozen at 24 hpi. For the control group, 200 zygotes were injected with 0.9 % NaCl. Total RNA was extracted using TRIzol reagent (Thermo Fisher Scientific) according to the manufacturer’s instructions. 500 ng of RNA was used as a template for cDNA synthesis using super script III reverse transcriptase (Thermo Fisher Scientific). Fast SYBR Green Master Mix (Thermo Fisher Scientific) was used for amplifying the cDNA. The samples were measured using the StepOnePlus Real-Time PCR System v2.3 (ABI, Thermo Fisher Scientific).

### GO functional analysis

Raw reads files obtained from Lewandowska et al. (2021) were aligned to *N. vectensis* reference genome (NCBI accession number: GCA_000209225.1) using STAR version 2.7.10a (Dobin, et al. 2013) and reads mapped to gene models were summarized using featureCounts Version 2.0.1 (Liao, et al. 2014). Differentially expressed genes between each treatment and control samples were identified using DESeq2 (Love, et al. 2014) and edgeR (Robinson, et al. 2010) (absolute log2 fold change > 1 and adjusted p value < 0.05). Only genes identified as differentially expressed by both the edgeR and DESeq2 methods were used in subsequent analyses (**Supplementary file 1**). We utilized blastp (Altschul, et al. 1990) and DIAMOND (Buchfink, et al. 2021) for functional annotation to identify the most similar homologs for each protein within the merged *N. vectensis* predicted proteome (comprising both JGI and Vienna annotations). These searches were performed against Uniprot database (UniProt 2023). Searching for ontology terms was carried out through QuickGO (Binns, et al. 2009). GO enrichment analysis was performed on sets of up-regulated genes using clusterProfiler (Wu, et al. 2021). A barplot per time point displayed up to 15 of the most significant GO terms (adjusted p-value < 0.05).

### Phylogenetic analysis

#### Sequence dataset construction

Putative metazoan caspases were identified by using tBLASTn and BLASTp searches with human caspases used as queries on NCBI, Uniprot and on downloaded genomes followed by reciprocal BLAST. Sequences with an e-value inferior to 1e-5 were retained. All identified sequences were analyzed with SMART (Letunic and Bork 2018) and InterProScan (EMBL-EBI) to verify the presence of caspase (CASc) domains. A full list of all protein sequences used in the BLAST search and phylogenetic analysis are provided in supplementary file 2. *Reticulomyxa filosa* (Foraminifiera) caspase homologues were used as outgroup. Protein sequence alignment was generated using the L-INS-i algorithm in MAFFT software version 7 (Katoh and Standley 2013), with default parameters.

#### Tree reconstruction

Phylogenetic analyses were conducted using amino-acid alignment (supplementary file 4), employing the Maximum-Likelihood (ML) method in IQ-TREE2 (Nguyen, et al. 2015) and Bayesian Inference (BI) analysis conducted in MrBayes version 3.2.6 (Ronquist, et al. 2012), respectively. Models of protein sequence evolution for ML (AIC criteria) and BI (BIC criteria) analyses were estimated using ModelFinder (Kalyaanamoorthy, et al. 2017). Branches support of the ML tree was assessed through 1000 ultrafast bootstrapping replicates (UBS) (Hoang, et al. 2018), 1000 bootstrap replicates for SH-aLRT (Guindon, et al. 2010) and an approximate aBayes test (Anisimova, et al. 2011). For bayesian inference analysis, two independent runs were carried out, each with four simultaneous Markov chains over 2,000,000 generations and sampled every 1000 generations. Convergence was checked using Tracer (Rambaut, et al. 2018). The consensus tree, along with the posterior probabilities of clades, was derived from trees sampled across the two runs. 25% of the sampled trees were discarded as burn-in. The resulting trees were visualized using figtree version 1.4.4 (<u>htp://tree.bio.ed.ac.uk/software/figtree/</u>).

#### Cross-species comparative transcriptomic analysis

To contextualize our findings within a comparative evolutionary framework, we analyzed the transcriptomic response to dsRNA viral mimic poly(I:C) across seven species (*N. vectensis*, *Homo sapiens*, *Mus musculus*, *S. kowalevskii*, *Crassostrea gigas*, *Branchiostoma belcheri*, and *Schmidtea mediterranea*) from diverse metazoan phylogenetic lineages. The aim was to ascertain whether dsRNA impacts apoptosis by examining the outcomes of the gene ontology (GO) analysis conducted in each study, and consequently, to identify if the genes within the apoptosis pathway are up-regulated. Detailed information on each of the studies used is found in supplementary file 3.

## References

Adamo A, Woglar A, Silva N, Penkner A, Jantsch V, La Volpe A. 2012. Transgene-mediated cosuppression and RNA interference enhance germ-line apoptosis in *Caenorhabditis elegans*. Proceedings of the National Academy of Sciences 109:3440–3445.

Admoni Y, Kozlovski I, Lewandowska M, Moran Y. 2020. TATA Binding Protein (TBP) Promoter Drives Ubiquitous Expression of Marker Transgene in the Adult Sea Anemone Nematostella vectensis. Genes 11:1081.

Alexopoulou L, Holt AC, Medzhitov R, Flavell RA. 2001. Recognition of double-stranded RNA and activation of NF-kappaB by Toll-like receptor 3. Nature 413:732–738.

Allocati N, Masulli M, Di Ilio C, De Laurenzi V. 2015. Die for the community: an overview of programmed cell death in bacteria. Cell death & disease 6:e1609–e1609.

Altschul SF, Gish W, Miller W, Myers EW, Lipman DJ. 1990. Basic local alignment search tool. J Mol Biol 215:403–410.

Anisimova M, Gil M, Dufayard JF, Dessimoz C, Gascuel O. 2011. Survey of branch support methods demonstrates accuracy, power, and robustness of fast likelihood-based approximation schemes. Syst Biol 60:685–699.

Bar Yaacov D. 2022. Functional analysis of ADARs in planarians supports a bilaterian ancestral role in suppressing double-stranded RNA-response. PLoS Pathog 18:e1010250.

Barber GN. 2001. Host defense, viruses and apoptosis. Cell Death Differ 8:113–126.

Barth ND, Subiros-Funosas R, Mendive-Tapia L, Duffin R, Shields MA, Cartwright JA, Henriques ST, Sot J, Goñi FM, Lavilla R. 2020. A fluorogenic cyclic peptide for imaging and quantification of drug-induced apoptosis. Nature communications 11:4027.

Baskić D, Popović S, Ristić P, Arsenijević NN. 2006. Analysis of cycloheximide-induced apoptosis in human leukocytes: Fluorescence microscopy using annexin V/propidium iodide versus acridin orange/ethidium bromide. Cell biology international 30:924–932.

Benedict CA, Norris PS, Ware CF. 2002. To kill or be killed: viral evasion of apoptosis. Nat Immunol 3:1013–1018.

Bergsbaken T, Fink SL, Cookson BT. 2009. Pyroptosis: host cell death and inflammation. Nature Reviews Microbiology 7:99–109.

Bertheloot D, Latz E, Franklin BS. 2021. Necroptosis, pyroptosis and apoptosis: an intricate game of cell death. Cell Mol Immunol 18:1106–1121.

Besch R, Poeck H, Hohenauer T, Senft D, Hacker G, Berking C, Hornung V, Endres S, Ruzicka T, Rothenfusser S, Hartmann G. 2009. Proapoptotic signaling induced by RIG-I and MDA-5 results in type I interferon-independent apoptosis in human melanoma cells. J Clin Invest 119:2399–2411.

Bidle KD, Falkowski PG. 2004. Cell death in planktonic, photosynthetic microorganisms. Nat Rev Microbiol 2:643–655.

Bieri T, Onishi M, Xiang T, Grossman AR, Pringle JR. 2016. Relative contributions of various cellular mechanisms to loss of algae during cnidarian bleaching. PLoS One 11:e0152693.

Binns D, Dimmer E, Huntley R, Barrell D, O’Donovan C, Apweiler R. 2009. QuickGO: a web-based tool for Gene Ontology searching. Bioinformatics 25:3045–3046.

Bouillet P, Metcalf D, Huang DC, Tarlinton DM, Kay TW, Kontgen F, Adams JM, Strasser A. 1999. Proapoptotic Bcl-2 relative Bim required for certain apoptotic responses, leukocyte homeostasis, and to preclude autoimmunity. Science 286:1735–1738.

Brennan JJ, Messerschmidt JL, Williams LM, Matthews BJ, Reynoso M, Gilmore TD. 2017. Sea anemone model has a single Toll-like receptor that can function in pathogen detection, NF-kappaB signal transduction, and development. Proc Natl Acad Sci U S A 114:E10122–E10131.

Buchfink B, Reuter K, Drost HG. 2021. Sensitive protein alignments at tree-of-life scale using DIAMOND. Nat Methods 18:366–368.

Carneiro BA, El-Deiry WS. 2020. Targeting apoptosis in cancer therapy. Nat Rev Clin Oncol 17:395–417.

Chan YK, Gack MU. 2016. Viral evasion of intracellular DNA and RNA sensing. Nature Reviews Microbiology 14:360–373.

Chattopadhyay S, Marques JT, Yamashita M, Peters KL, Smith K, Desai A, Williams BR, Sen GC. 2010a. Viral apoptosis is induced by IRF-3-mediated activation of Bax. EMBO J 29:1762–1773.

Chattopadhyay S, Marques JT, Yamashita M, Peters KL, Smith K, Desai A, Williams BR, Sen GC. 2010b. Viral apoptosis is induced by IRF-3-mediated activation of Bax. The EMBO journal 29:1762–1773.

Chen S, Li S, Chen H, Gong Y, Yang D, Zhang Y, Liu Q. 2023a. Caspase-mediated LPS sensing and pyroptosis signaling in Hydra. Science Advances 9:eadh4054.

Chen S, Li S, Chen H, Gong Y, Yang D, Zhang Y, Liu Q. 2023b. Caspase-mediated LPS sensing and pyroptosis signaling in Hydra. Sci Adv 9:eadh4054.

Chen YG, Hur S. 2022. Cellular origins of dsRNA, their recognition and consequences. Nature Reviews Molecular Cell Biology 23:286–301.

Crowley LC, Marfell BJ, Scott AP, Waterhouse NJ. 2016. Quantitation of apoptosis and necrosis by annexin V binding, propidium iodide uptake, and flow cytometry. Cold Spring Harb Protoc 2016:953–957.

David CN, Schmidt N, Schade M, Pauly B, Alexandrova O, Bottger A. 2005. Hydra and the evolution of apoptosis. Integr Comp Biol 45:631–638.

Deponte M. 2008. Programmed cell death in protists. Biochim Biophys Acta 1783:1396–1405.

Dixit E, Kagan JC. 2013. Intracellular pathogen detection by RIG-I-like receptors. Adv Immunol 117:99–125.

Dobin A, Davis CA, Schlesinger F, Drenkow J, Zaleski C, Jha S, Batut P, Chaisson M, Gingeras TR. 2013. STAR: ultrafast universal RNA-seq aligner. Bioinformatics 29:15–21.

El Maadidi S, Faletti L, Berg B, Wenzl C, Wieland K, Chen ZJ, Maurer U, Borner C. 2014. A novel mitochondrial MAVS/Caspase-8 platform links RNA virus-induced innate antiviral signaling to Bax/Bak-independent apoptosis. J Immunol 192:1171–1183.

Erwin DH, Laflamme M, Tweedt SM, Sperling EA, Pisani D, Peterson KJ. 2011. The Cambrian conundrum: early divergence and later ecological success in the early history of animals. Science 334:1091–1097.

Estornes Y, Toscano F, Virard F, Jacquemin G, Pierrot A, Vanbervliet B, Bonnin M, Lalaoui N, Mercier-Gouy P, Pachéco Y, et al. 2012. dsRNA induces apoptosis through an atypical death complex associating TLR3 to caspase-8. Cell Death && Differentiation 19:1482–1494.

Genikhovich G, Technau U. 2009. Induction of spawning in the starlet sea anemone Nematostella vectensis, in vitro fertilization of gametes, and dejellying of zygotes. Cold Spring Harbor Protocols 2009:pdb. prot5281.

Gitlin L, Barchet W, Gilfillan S, Cella M, Beutler B, Flavell RA, Diamond MS, Colonna M. 2006. Essential role of mda-5 in type I IFN responses to polyriboinosinic:polyribocytidylic acid and encephalomyocarditis picornavirus. Proc Natl Acad Sci U S A 103:8459–8464.

Glas M, Coch C, Trageser D, Dassler J, Simon M, Koch P, Mertens J, Quandel T, Gorris R, Reinartz R, et al. 2013. Targeting the cytosolic innate immune receptors RIG-I and MDA5 effectively counteracts cancer cell heterogeneity in glioblastoma. Stem Cells 31:1064–1074.

Goubau D, Schlee M, Deddouche S, Pruijssers AJ, Zillinger T, Goldeck M, Schuberth C, Van der Veen AG, Fujimura T, Rehwinkel J, et al. 2014. Antiviral immunity via RIG-I-mediated recognition of RNA bearing 5’-diphosphates. Nature 514:372–375.

Guindon S, Dufayard JF, Lefort V, Anisimova M, Hordijk W, Gascuel O. 2010. New algorithms and methods to estimate maximum-likelihood phylogenies: assessing the performance of PhyML 3.0. Syst Biol 59:307–321.

Gurtu V, Kain SR, Zhang G. 1997. Fluorometric and colorimetric detection of caspase activity associated with apoptosis. Anal Biochem 251:98–102.

Hagai T, Chen X, Miragaia RJ, Rostom R, Gomes T, Kunowska N, Henriksson J, Park JE, Proserpio V, Donati G, et al. 2018. Gene expression variability across cells and species shapes innate immunity. Nature 563:197–202.

Hamann A, Brust D, Osiewacz HD. 2008. Apoptosis pathways in fungal growth, development and ageing. Trends Microbiol 16:276–283.

Hoang DT, Chernomor O, von Haeseler A, Minh BQ, Vinh LS. 2018. UFBoot2: Improving the Ultrafast Bootstrap Approximation. Mol Biol Evol 35:518–522.

Ivashkiv LB, Donlin LT. 2014. Regulation of type I interferon responses. Nat Rev Immunol 14:36–49.

Iwama RE, Moran Y. 2023. Origins and diversification of animal innate immune responses against viral infections. Nat Ecol Evol 7:182–193.

Jiang S, Zhou Z, Sun Y, Zhang T, Sun L. 2020. Coral gasdermin triggers pyroptosis. Science Immunology 5:eabd2591.

Johnson AG, Wein T, Mayer ML, Duncan-Lowey B, Yirmiya E, Oppenheimer-Shaanan Y, Amitai G, Sorek R, Kranzusch PJ. 2022. Bacterial gasdermins reveal an ancient mechanism of cell death. Science 375:221–225.

Kaczanowski S, Sajid M, Reece SE. 2011. Evolution of apoptosis-like programmed cell death in unicellular protozoan parasites. Parasit Vectors 4:44.

Kalyaanamoorthy S, Minh BQ, Wong TKF, von Haeseler A, Jermiin LS. 2017. ModelFinder: fast model selection for accurate phylogenetic estimates. Nat Methods 14:587–589.

Kato H, Takeuchi O, Mikamo-Satoh E, Hirai R, Kawai T, Matsushita K, Hiiragi A, Dermody TS, Fujita T, Akira S. 2008. Length-dependent recognition of double-stranded ribonucleic acids by retinoic acid-inducible gene-I and melanoma differentiation-associated gene 5. J Exp Med 205:1601–1610.

Kato H, Takeuchi O, Sato S, Yoneyama M, Yamamoto M, Matsui K, Uematsu S, Jung A, Kawai T, Ishii KJ, et al. 2006. Differential roles of MDA5 and RIG-I helicases in the recognition of RNA viruses. Nature 441:101–105.

Katoh K, Standley DM. 2013. MAFFT multiple sequence alignment software version 7: improvements in performance and usability. Mol Biol Evol 30:772–780.

Koonin E, Aravind L. 2002a. Origin and evolution of eukaryotic apoptosis: the bacterial connection. Cell Death & Differentiation 9:394–404.

Koonin EV, Aravind L. 2002b. Origin and evolution of eukaryotic apoptosis: the bacterial connection. Cell Death Differ 9:394–404.

Krasovec G, Horkan HR, Quéinnec É, Chambon J-P. 2023. Intrinsic apoptosis is evolutionarily divergent among metazoans. Evolution Letters.

Kumar S, Ingle H, Mishra S, Mahla RS, Kumar A, Kawai T, Akira S, Takaoka A, Raut AA, Kumar H. 2015. IPS-1 differentially induces TRAIL, BCL2, BIRC3 and PRKCE in type I interferons-dependent and -independent anticancer activity. Cell Death Dis 6:e1758.

Kvitt H, Kramarsky-Winter E, Maor-Landaw K, Zandbank K, Kushmaro A, Rosenfeld H, Fine M, Tchernov D. 2015. Breakdown of coral colonial form under reduced pH conditions is initiated in polyps and mediated through apoptosis. Proceedings of the National Academy of Sciences 112:2082–2086.

Lafont M, Vergnes A, Vidal-Dupiol J, de Lorgeril J, Gueguen Y, Haffner P, Petton B, Chaparro C, Barrachina C, Destoumieux-Garzon D, et al. 2020. A Sustained Immune Response Supports Long-Term Antiviral Immune Priming in the Pacific Oyster, Crassostrea gigas. mBio 11.

Lamkanfi M, Declercq W, Kalai M, Saelens X, Vandenabeele P. 2002. Alice in caspase land. A phylogenetic analysis of caspases from worm to man. Cell Death Differ 9:358–361.

Lasi M, Pauly B, Schmidt N, Cikala M, Stiening B, Kasbauer T, Zenner G, Popp T, Wagner A, Knapp RT, et al. 2010. The molecular cell death machinery in the simple cnidarian Hydra includes an expanded caspase family and pro- and anti-apoptotic Bcl-2 proteins. Cell Res 20:812–825.

Layden MJ, Rentzsch F, Röttinger E. 2016. The rise of the starlet sea anemone Nematostella vectensis as a model system to investigate development and regeneration. Wiley Interdisciplinary Reviews: Developmental Biology 5:408–428.

Letunic I, Bork P. 2018. 20 years of the SMART protein domain annotation resource. Nucleic Acids Res 46:D493–D496.

Lewandowska M, Sharoni T, Admoni Y, Aharoni R, Moran Y. 2021. Functional Characterization of the Cnidarian Antiviral Immune Response Reveals Ancestral Complexity. Mol Biol Evol 38:4546–4561.

Li J, Yuan J. 2008. Caspases in apoptosis and beyond. Oncogene 27:6194–6206.

Li P, Nijhawan D, Budihardjo I, Srinivasula SM, Ahmad M, Alnemri ES, Wang X. 1997. Cytochrome c and dATP-dependent formation of Apaf-1/caspase-9 complex initiates an apoptotic protease cascade. Cell 91:479–489.

Liao Y, Smyth GK, Shi W. 2014. featureCounts: an efficient general purpose program for assigning sequence reads to genomic features. Bioinformatics 30:923–930.

Linehan MM, Dickey TH, Molinari ES, Fitzgerald ME, Potapova O, Iwasaki A, Pyle AM. 2018. A minimal RNA ligand for potent RIG-I activation in living mice. Sci Adv 4:e1701854.

Love MI, Huber W, Anders S. 2014. Moderated estimation of fold change and dispersion for RNA-seq data with DESeq2. Genome Biol 15:550.

Madeo F, Herker E, Maldener C, Wissing S, Lachelt S, Herlan M, Fehr M, Lauber K, Sigrist SJ, Wesselborg S, Frohlich KU. 2002. A caspase-related protease regulates apoptosis in yeast. Mol Cell 9:911–917.

Matsumoto M, Seya T. 2008. TLR3: interferon induction by double-stranded RNA including poly(I:C). Adv Drug Deliv Rev 60:805–812.

Nakamura H, Yoshida M, Tsuiki H, Ito K, Ueno M, Nakao M, Oka K, Tada M, Kochi M, Kuratsu J, et al. 1998. Identification of a human homolog of the Drosophila neuralized gene within the 10q25.1 malignant astrocytoma deletion region. Oncogene 16:1009–1019.

Nakatsumi H, Yonehara S. 2010. Identification of functional regions defining different activity in caspase-3 and caspase-7 within cells. J Biol Chem 285:25418–25425.

Nguyen LT, Schmidt HA, von Haeseler A, Minh BQ. 2015. IQ-TREE: a fast and effective stochastic algorithm for estimating maximum-likelihood phylogenies. Mol Biol Evol 32:268–274.

Nicholson DW, Ali A, Thornberry NA, Vaillancourt JP, Ding CK, Gallant M, Gareau Y, Griffin PR, Labelle M, Lazebnik YA, et al. 1995. Identification and inhibition of the ICE/CED-3 protease necessary for mammalian apoptosis. Nature 376:37–43.

Oberst A, Bender C, Green DR. 2008. Living with death: the evolution of the mitochondrial pathway of apoptosis in animals. Cell Death Differ 15:1139–1146.

Orzalli MH, Kagan JC. 2017. Apoptosis and necroptosis as host defense strategies to prevent viral infection. Trends in cell biology 27:800–809.

Pankow S, Bamberger C. 2007. The p53 tumor suppressor-like protein nvp63 mediates selective germ cell death in the sea anemone Nematostella vectensis. PLoS One 2:e782.

Pasparakis M, Vandenabeele P. 2015. Necroptosis and its role in inflammation. Nature 517:311–320.

Pernice M, Dunn SR, Miard T, Dufour S, Dove S, Hoegh-Guldberg O. 2011. Regulation of apoptotic mediators reveals dynamic responses to thermal stress in the reef building coral Acropora millepora. PLoS One 6:e16095.

Pirnia F, Schneider E, Betticher DC, Borner MM. 2002. Mitomycin C induces apoptosis and caspase-8 and-9 processing through a caspase-3 and Fas-independent pathway. Cell Death && Differentiation 9:905–914.

Pollitt LC, Colegrave N, Khan SM, Sajid M, Reece SE. 2010. Investigating the evolution of apoptosis in malaria parasites: the importance of ecology. Parasit Vectors 3:105.

Quistad SD, Stotland A, Barott KL, Smurthwaite CA, Hilton BJ, Grasis JA, Wolkowicz R, Rohwer FL. 2014. Evolution of TNF-induced apoptosis reveals 550 My of functional conservation. Proceedings of the National Academy of Sciences 111:9567–9572.

Rambaut A, Drummond AJ, Xie D, Baele G, Suchard MA. 2018. Posterior Summarization in Bayesian Phylogenetics Using Tracer 1.7. Syst Biol 67:901–904.

Reape TJ, McCabe PF. 2010. Apoptotic-like regulation of programmed cell death in plants. Apoptosis 15:249–256.

Rehwinkel J, Gack MU. 2020. RIG-I-like receptors: their regulation and roles in RNA sensing. Nat Rev Immunol 20:537–551.

Robinson MD, McCarthy DJ, Smyth GK. 2010. edgeR: a Bioconductor package for differential expression analysis of digital gene expression data. Bioinformatics 26:139–140.

Ronquist F, Teslenko M, van der Mark P, Ayres DL, Darling A, Hohna S, Larget B, Liu L, Suchard MA, Huelsenbeck JP. 2012. MrBayes 3.2: efficient Bayesian phylogenetic inference and model choice across a large model space. Syst Biol 61:539–542.

Ros M, Pernice M, Le Guillou S, Doblin MA, Schrameyer V, Laczka O. 2016. Colorimetric detection of caspase 3 activity and reactive oxygen derivatives: Potential early indicators of thermal stress in corals. Journal of Marine Sciences 2016.

Rottinger E. 2021. Nematostella vectensis, an Emerging Model for Deciphering the Molecular and Cellular Mechanisms Underlying Whole-Body Regeneration. Cells 10.

Rousset F, Sorek R. 2023. The evolutionary success of regulated cell death in bacterial immunity. Curr Opin Microbiol 74:102312.

Sakamaki K, Satou Y. 2009. Caspases: evolutionary aspects of their functions in vertebrates. J Fish Biol 74:727–753.

Schlee M, Hartmann G. 2016. Discriminating self from non-self in nucleic acid sensing. Nature Reviews Immunology 16:566–580.

Seipp S, Schmich J, Leitz T. 2001. Apoptosis--a death-inducing mechanism tightly linked with morphogenesis in Hydractina echinata (Cnidaria, Hydrozoa). Development 128:4891–4898.

Sharon A, Finkelstein A, Shlezinger N, Hatam I. 2009. Fungal apoptosis: function, genes and gene function. FEMS Microbiol Rev 33:833–854.

Son KN, Liang Z, Lipton HL. 2015. Double-Stranded RNA Is Detected by Immunofluorescence Analysis in RNA and DNA Virus Infections, Including Those by Negative-Stranded RNA Viruses. J Virol 89:9383–9392.

Stetson DB, Medzhitov R. 2006. Type I interferons in host defense. Immunity 25:373–381.

Sun Q, Sun L, Liu HH, Chen X, Seth RB, Forman J, Chen ZJ. 2006. The specific and essential role of MAVS in antiviral innate immune responses. Immunity 24:633–642.

Tait SW, Green DR. 2010. Mitochondria and cell death: outer membrane permeabilization and beyond. Nature Reviews Molecular Cell Biology 11:621–632.

Talice S, Barkan SK, Snyder GA, Ottolenghi A, Eliachar S, Ben-Romano R, Oisher S, Sharoni T, Lewandowska M, Sultan E, et al. 2023. Candidate Stem Cell Isolation and Transplantation in Hexacorallia. In: Elsevier BV.

Tassia MG, Hallowell HA, Waits DS, Range RC, Lowe CJ, Graze RM, Schwartz EH, Halanych KM. 2023. Induced Immune Reaction in the Acorn Worm, Saccoglossus kowalevskii, Informs the Evolution of Antiviral Immunity. Mol Biol Evol 40.

Taylor RC, Cullen SP, Martin SJ. 2008. Apoptosis: controlled demolition at the cellular level. Nature Reviews Molecular Cell Biology 9:231–241.

Tchernov D, Kvitt H, Haramaty L, Bibby TS, Gorbunov MY, Rosenfeld H, Falkowski PG. 2011. Apoptosis and the selective survival of host animals following thermal bleaching in zooxanthellate corals. Proceedings of the National Academy of Sciences 108:9905–9909.

Technau U, Steele RE. 2011. Evolutionary crossroads in developmental biology: Cnidaria. Development 138:1447–1458.

Thomsen ND, Koerber JT, Wells JA. 2013. Structural snapshots reveal distinct mechanisms of procaspase-3 and -7 activation. Proc Natl Acad Sci U S A 110:8477–8482.

Tomasz M. 1995. Mitomycin C: small, fast and deadly (but very selective). Chem Biol 2:575–579.

UniProt C. 2023. UniProt: the Universal Protein Knowledgebase in 2023. Nucleic Acids Res 51:D523–D531.

Upton JW, Chan FK. 2014. Staying alive: cell death in antiviral immunity. Mol Cell 54:273–280.

van Dommelen SL, Sumaria N, Schreiber RD, Scalzo AA, Smyth MJ, Degli-Esposti MA. 2006. Perforin and granzymes have distinct roles in defensive immunity and immunopathology. Immunity 25:835–848.

Verdonck S, Nemegeer J, Vandenabeele P, Maelfait J. 2022. Viral manipulation of host cell necroptosis and pyroptosis. Trends Microbiol 30:593–605.

Walsh JG, Cullen SP, Sheridan C, Luthi AU, Gerner C, Martin SJ. 2008. Executioner caspase-3 and caspase-7 are functionally distinct proteases. Proc Natl Acad Sci U S A 105:12815–12819.

Weber F, Wagner V, Rasmussen SB, Hartmann R, Paludan SR. 2006. Double-stranded RNA is produced by positive-strand RNA viruses and DNA viruses but not in detectable amounts by negative-strand RNA viruses. J Virol 80:5059–5064.

Wiens M, Krasko A, Blumbach B, Müller I, Müller W. 2000. Increased expression of the potential proapoptotic molecule DD2 and increased synthesis of leukotriene B4 during allograft rejection in a marine sponge. Cell Death & Differentiation 7:461–469.

Wiens M, Krasko A, Müller CI, Müller WE. 2000. Molecular evolution of apoptotic pathways: cloning of key domains from sponges (Bcl-2 homology domains and death domains) and their phylogenetic relationships. Journal of Molecular Evolution 50:520–531.

Wu T, Hu E, Xu S, Chen M, Guo P, Dai Z, Feng T, Zhou L, Tang W, Zhan L, et al. 2021. clusterProfiler 4.0: A universal enrichment tool for interpreting omics data. Innovation (Camb) 2:100141.

Xia S, Hollingsworth LRt, Wu H. 2020. Mechanism and Regulation of Gasdermin-Mediated Cell Death. Cold Spring Harb Perspect Biol 12.

Yoneyama M, Kikuchi M, Natsukawa T, Shinobu N, Imaizumi T, Miyagishi M, Taira K, Akira S, Fujita T. 2004. The RNA helicase RIG-I has an essential function in double-stranded RNA-induced innate antiviral responses. Nat Immunol 5:730–737.

Yuan J, Horvitz HR. 2004. A first insight into the molecular mechanisms of apoptosis. Cell 116:S53–56.

Yuan J, Shaham S, Ledoux S, Ellis HM, Horvitz HR. 1993. The C. elegans cell death gene ced-3 encodes a protein similar to mammalian interleukin-1β-converting enzyme. Cell 75:641–652.

Zhang QL, Qiu HY, Liang MZ, Luo B, Wang XQ, Chen JY. 2017. Exploring gene expression changes in the amphioxus gill after poly(I:C) challenge using digital expression profiling. Fish Shellfish Immunol 70:57–65.

