## Supplementary material for "Induction of apoptosis by double-stranded RNA was present in the last common ancestor of cnidarian and bilaterian animals": Figure S1

a

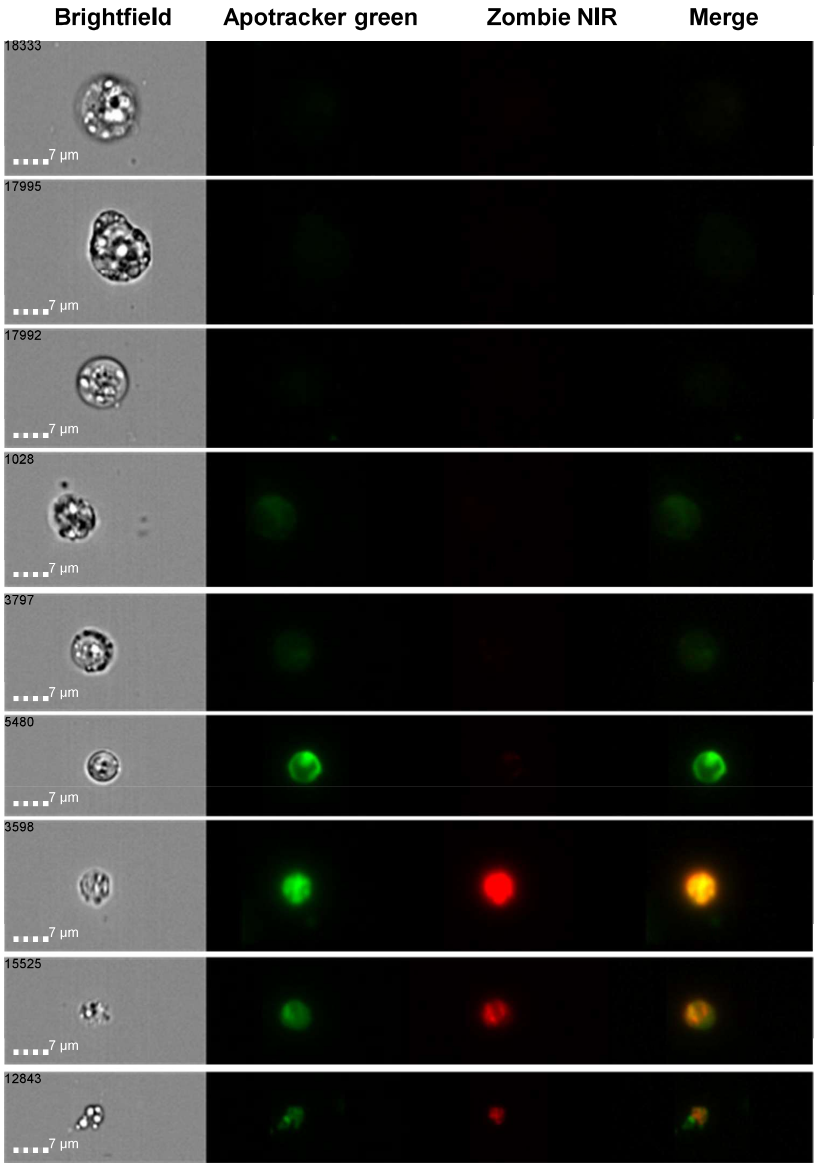


b

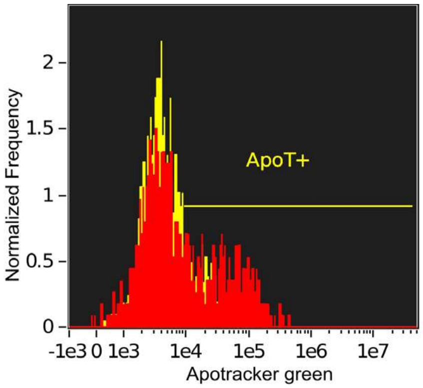


Intensity_Apotracker green

| Population | Count | %Gated | Mean | Median | CV |
|---|---|---|---|---|---|
| CHX whole population | 1133 | 100 | 24231.08 | 5502.5 | 188.75 |
| CHX ApoT+ | 412 | 36.4 | 59869.57 | 37658.24 | 102.32 |
| DMSO whole population | 2073 | 100 | 9567.12 | 4360.35 | 198.99 |
| DMSO ApoT+ | 413 | 19.9 | 31366.02 | 20673.04 | 110.94 |

c

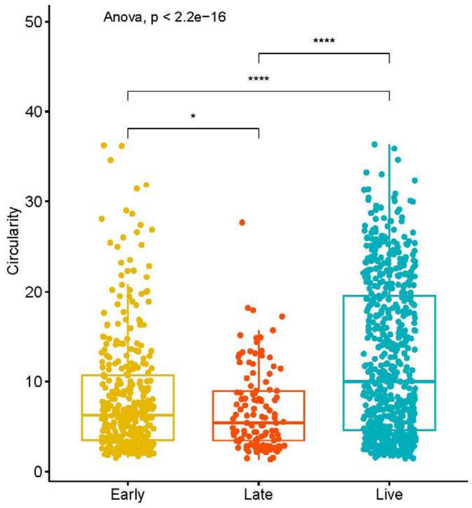
