## Supplementary figures and images for "Induction of apoptosis by double-stranded RNA was present in the last common ancestor of cnidarian and bilaterian animals"

### Figure S2

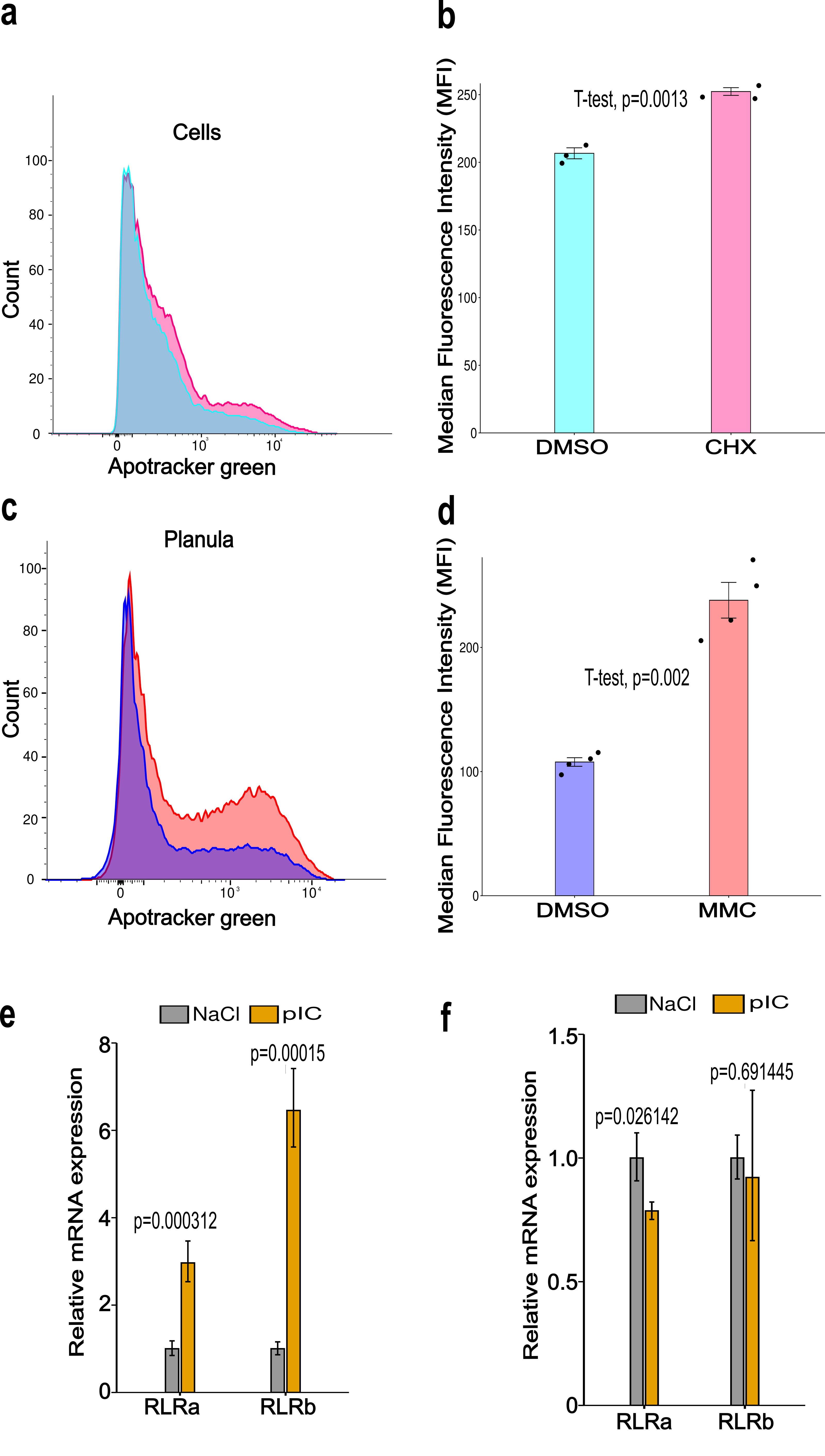

### Figure S3

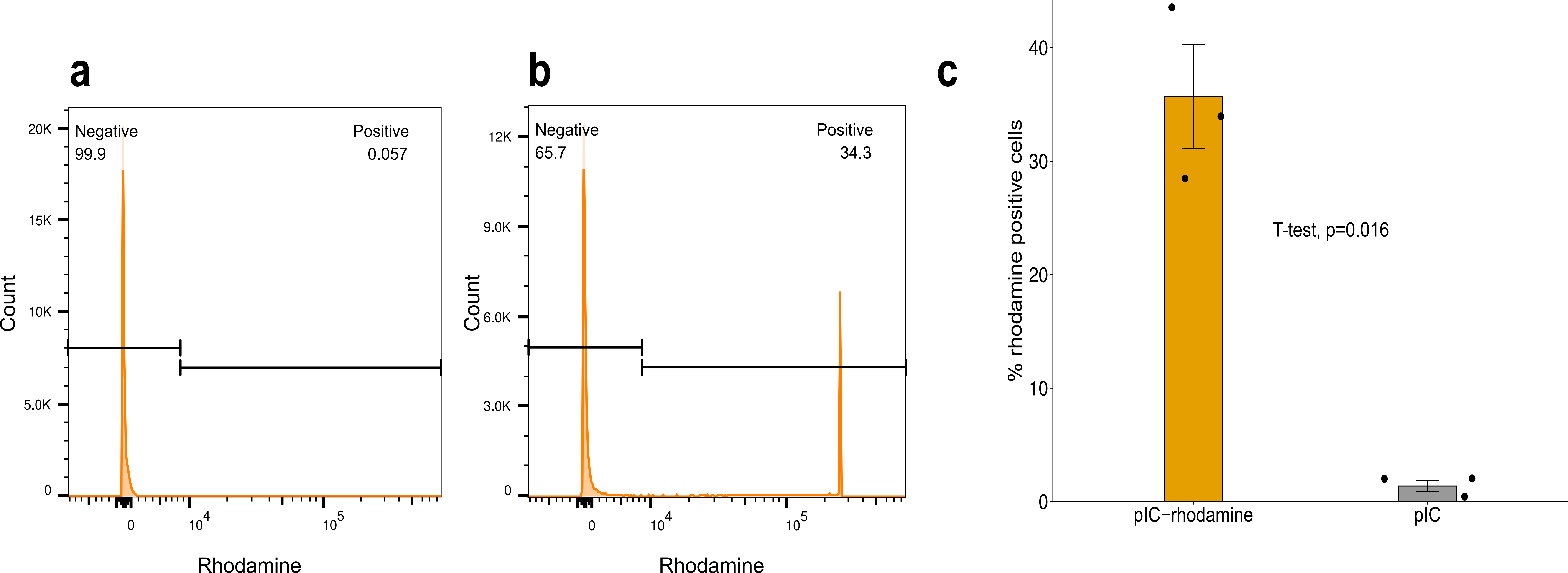

### Figure S4

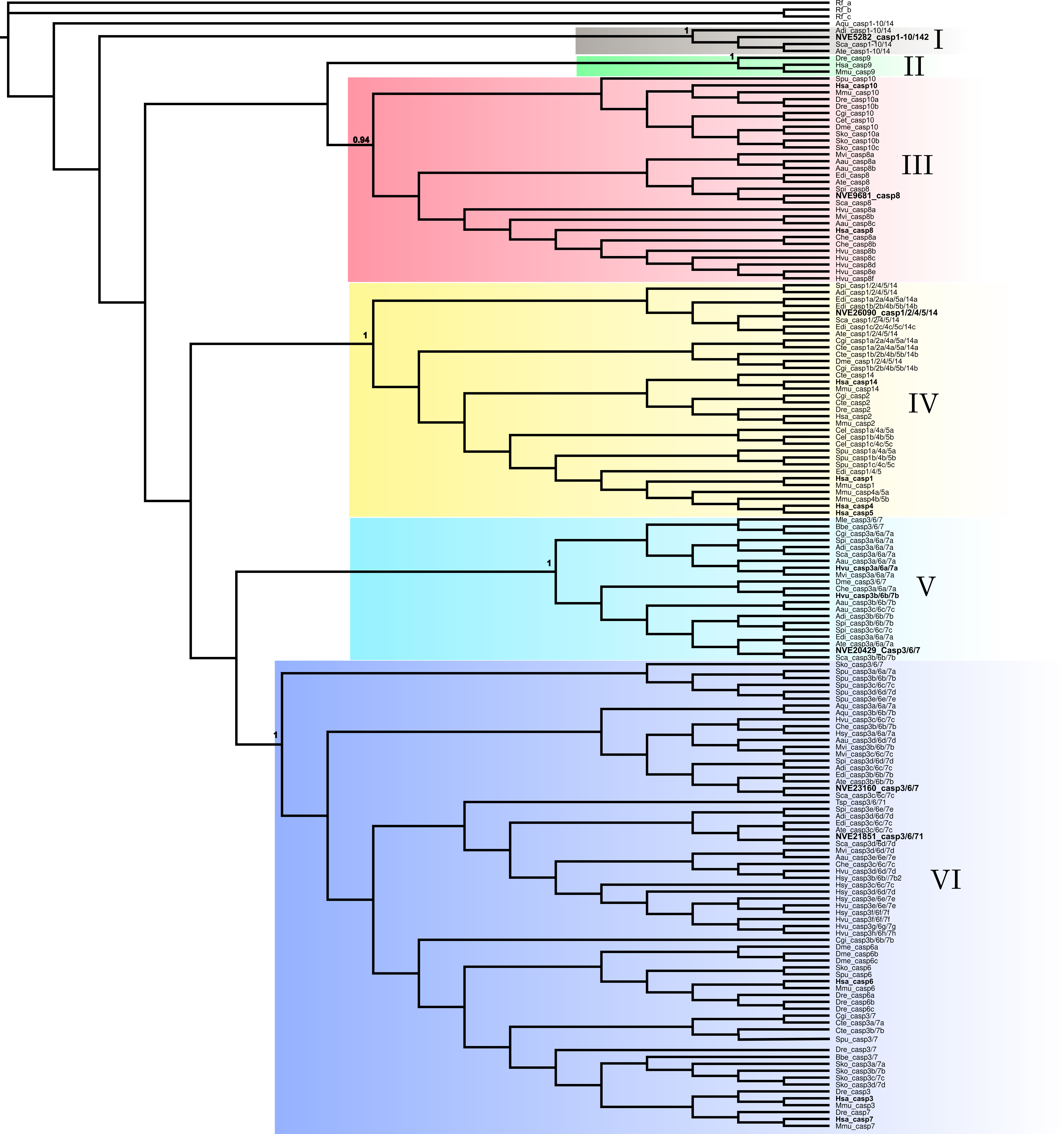
