## Supplementary File 2 for "Induction of apoptosis by double-stranded RNA was present in the last common ancestor of cnidarian and bilaterian animals"

**Supplementary file 2:** Accession numbers for caspase protein sequences used in the BLAST search and phylogenetic analysis.

| **Name** | **Species** | **Genbank ID/Uniprot ID** |
| --- | --- | --- |
| Rfi_a | *Reticulomyxa filosa* | X6N4E9 |
| Rfi_b | *Reticulomyxa filosa* | X6NIS5 |
| Rfi_c | *Reticulomyxa filosa* | X6ME08 |
| Mle_casp3/6/7 | *Mnemiopsis leidyi* | GCA_000226015.1 |
| Aqu_casp3a/6a/7a | *Amphimedon queenslandica* | A0A1X7VNG6 |
| Aqu_casp3b/6b/7b | *Amphimedon queenslandica* | A0A1X7UIG0 |
| Aqu_casp1-10/14 | *Amphimedon queenslandica* | GCA_000090795.2 |
| Tsp_casp3/6/7 | *Trichoplax_sp* | A0A369SE24 |
| NVE23160_casp3/6/7 | *Nematostella vectensis* | GCF_932526225.1 |
| NVE9681_casp8 | *Nematostella vectensis* | GCF_932526225.1 |
| NVE5282_casp1-10/14 | *Nematostella vectensis* | GCF_932526225.1 |
| NVE20429_casp3/6/7 | *Nematostella vectensis* | GCF_932526225.1 |
| NVE21851casp3/6/7 | *Nematostella vectensis* | GCF_932526225.1 |
| NVE26090_casp1/2/4/5/14 | *Nematostella vectensis* | GCF_932526225.1 |
| Sca_casp3a/6a/7a | *Scolanthus callimorphis* | <https://simrbase.stowers.org/wormanemone> |
| Sca_casp3c/6c/7c | *Scolanthus callimorphis* | <https://simrbase.stowers.org/wormanemone> |
| Sca_casp1-10/14 | *Scolanthus callimorphis* | <https://simrbase.stowers.org/wormanemone> |
| Sca_casp3b/6b/7b | *Scolanthus callimorphis* | <https://simrbase.stowers.org/wormanemone> |
| Sca_casp3d/6d/7d | *Scolanthus callimorphis* | <https://simrbase.stowers.org/wormanemone> |
| Sca_casp8 | *Scolanthus callimorphis* | <https://simrbase.stowers.org/wormanemone> |
| Sca_casp1/2/4/5/14 | *Scolanthus callimorphis* | <https://simrbase.stowers.org/wormanemone> |
| Ate_casp8 | *Actinia tenebrosa* | GCF_009602425.1 |
| Ate_casp3b/6b/7b | *Actinia tenebrosa* | GCF_009602425.1 |
| Ate_casp1-10/14 | *Actinia tenebrosa* | GCF_009602425.1 |
| Ate_casp1/2/3/5/14 | *Actinia tenebrosa* | GCF_009602425.1 |
| Ate_casp3c/6c/7c | *Actinia tenebrosa* | GCF_009602425.1 |
| Ate_casp3a/6a/7a | *Actinia tenebrosa* | GCF_009602425.1 |
| Spi_casp3d/6d/7d | *Stylophora pistillata* | A0A2B4RVA9 |
| Spi_casp3b/6b/7b | *Stylophora pistillata* | A0A2B4SJP8 |
| Spi_casp3a/6a/7a | *Stylophora pistillata* | A0A2B4RBN1 |
| Spi_casp8 | *Stylophora pistillata* | A0A2B4RVR5 |
| Spi_casp1/2/4/5/14 | *Stylophora pistillata* | A0A2B4SCW1 |
| Spi_casp3e/6e/7e | *Stylophora pistillata* | A0A2B4SWS7 |
| Spi_casp3c/6c/7c | *Stylophora pistillata* | A0A2B4SF88 |
| Adi_casp3d/6d/7d | *Acropora digitifera* | GCA_000222465.2 |
| Adi_casp3c/6c/7c | *Acropora digitifera* | GCA_000222465.2 |
| Adi_casp3a/6a/7a | *Acropora digitifera* | GCA_000222465.2 |
| Adi_casp1-10/14 | *Acropora digitifera* | GCA_000222465.2 |
| Adi_casp3b/6b/7b | *Acropora digitifera* | GCA_000222465.2 |
| Adi_casp1/2/4/5/14 | *Acropora digitifera* | GCA_000222465.2 |
| Edi_casp3b/6b/7b | *Exaiptasia diaphana* | GCA_001417965.1 |
| Edi_casp1c/2c/4c/5c/14c | *Exaiptasia diaphana* | GCA_001417965.1 |
| Edi_casp8 | *Exaiptasia diaphana* | GCA_001417965.1 |
| Edi_casp3c/6c/7c | *Exaiptasia diaphana* | GCA_001417965.1 |
| Edi_casp3a/6a/7a | *Exaiptasia diaphana* | GCA_001417965.1 |
| Edi_casp1b/2b/4b/5b/14b | *Exaiptasia diaphana* | GCA_001417965.1 |
| Edi_casp1a/2a/4a/5a/14a | *Exaiptasia diaphana* | GCA_001417965.1 |
| Edi_casp1/4/5 | *Exaiptasia diaphana* | GCA_001417965.1 |
| Che_casp3b/6b/7b | *Clytia hemisphaerica* | GCA_902728285.1 |
| Che_casp3a/6a/7a | *Clytia hemisphaerica* | GCA_902728285.1 |
| Che_casp8a | *Clytia hemisphaerica* | GCA_902728285.1 |
| Che_casp3c/6c/7c | *Clytia hemisphaerica* | GCA_902728285.1 |
| Che_casp8b | *Clytia hemisphaerica* | GCA_902728285.1 |
| Hvu_casp3c/6c/7c | *Hydra vulgaris* | D1MAR4 |
| Hvu_casp3b/6b/7b | *Hydra vulgaris* | Q9GV89 |
| Hvu_casp3e/6e/7e | *Hydra vulgaris* | A0A8B6XP53 |
| Hvu_casp3h/6h/7h | *Hydra vulgaris* | A0A8B6XQI4 |
| Hvu_casp3d/6d/7d | *Hydra vulgaris* | E2DGP9 |
| Hvu_casp3f/6f/7f | *Hydra vulgaris* | A0A8B6XIS7 |
| Hvu_casp8c | *Hydra vulgaris* | XP_047129731.1 |
| Hvu_casp8e | *Hydra vulgaris* | A0A8B7DUV5 |
| Hvu_casp8b | *Hydra vulgaris* | A0A8B7DEI2 |
| Hvu_casp8f | *Hydra vulgaris* | A0A8B7DAF8 |
| Hvu_casp8d | *Hydra vulgaris* | XP_047134542.1 |
| Hvu_casp3g/6g/7g | *Hydra vulgaris* | XP_047134596.1 |
| Hvu_casp3a/6a/7a | *Hydra vulgaris* | NP_001296709.1 |
| Hvu_casp8a | *Hydra vulgaris* | NP_001267753.1 |
| Hsy_casp3a/6a/7a | *Hydractinia symbiolongicarpus* | GCA_029227915.2 |
| Hsy_casp3b/6b//7b | *Hydractinia symbiolongicarpus* | GCA_029227915.2 |
| Hsy_casp3d/6d/7d | *Hydractinia symbiolongicarpus* | GCA_029227915.2 |
| Hsy_casp3f/6f/7f | *Hydractinia symbiolongicarpus* | GCA_029227915.2 |
| Hsy_casp3c/6c/7c | *Hydractinia symbiolongicarpus* | GCA_029227915.2 |
| Hsy_casp3e/6e/7e | *Hydractinia symbiolongicarpus* | GCA_029227915.2 |
| Mvi_casp3b/6b/7b | *Morbakka virulenta* | GCA_003991215.1 |
| Mvi_casp8b | *Morbakka virulenta* | GCA_003991215.1 |
| Mvi_casp3d/6d/7d | *Morbakka virulenta* | GCA_003991215.1 |
| Mvi_casp3c/6c/7c | *Morbakka virulenta* | GCA_003991215.1 |
| Mvi_casp8a | *Morbakka virulenta* | GCA_003991215.1 |
| Mvi_casp3a/6a/7a | *Morbakka virulenta* | GCA_003991215.1 |
| Aau_casp3b/6b/7b | *Aurelia aurita* | GCA_004194415.1 |
| Aau_casp8a | *Aurelia aurita* | GCA_004194415.1 |
| Aau_casp8b | *Aurelia aurita* | GCA_004194415.1 |
| Aau_casp3c/6c/7c | *Aurelia aurita* | GCA_004194415.1 |
| Aau_casp3d/6d/7d | *Aurelia aurita* | GCA_004194415.1 |
| Aau_casp3e/6e/7e | *Aurelia aurita* | GCA_004194415.1 |
| Aau_casp3a/6a/7a | *Aurelia aurita* | GCA_004194415.1 |
| Aau_casp8c | *Aurelia aurita* | GCA_004194415.1 |
| Cgi_casp2 | *Crassostrea gigas* | GCA_902806645.1 |
| Cgi_casp1a/2a/4a/5a/14a | *Crassostrea gigas* | GCA_902806645.1 |
| Cgi_casp10 | *Crassostrea gigas* | GCA_902806645.1 |
| Cgi_casp1b/2b/4b/5b/14b | *Crassostrea gigas* | GCA_902806645.1 |
| Cgi_casp3a/6a/7a | *Crassostrea gigas* | GCA_902806645.1 |
| Cgi_casp3b/6b/7b | *Crassostrea gigas* | GCA_902806645.1 |
| Cgi_casp3/7 | *Crassostrea gigas* | GCA_902806645.1 |
| Cte_casp1a/2a/4a/5a/14a | *Capitella teleta* | GCA_000328365.1 |
| Cte_casp3b/7b | *Capitella teleta* | GCA_000328365.1 |
| Cte_casp2 | *Capitella teleta* | GCA_000328365.1 |
| Cte_casp10 | *Capitella teleta* | GCA_000328365.1 |
| Cte_casp1b/2b/4b/5b/14b | *Capitella teleta* | GCA_000328365.1 |
| Cte_casp14 | *Capitella teleta* | GCA_000328365.1 |
| Cte_casp3a/7a | *Capitella teleta* | GCA_000328365.1 |
| Dme_casp1/2/4/5/14 | *Drosophila melanogaster* | Q9XYF4 |
| Dme_casp10 | *Drosophila melanogaster* | Q8IRY7 |
| Dme_casp3/6/7 | *Drosophila melanogaster* | Q7KHI6 |
| Dme_casp6b | *Drosophila melanogaster* | O02002 |
| Dme_casp6a | *Drosophila melanogaster* | Q9VET9 |
| Dme_casp6c | *Drosophila melanogaster* | O01382 |
| Cel_casp1a/4a/5a | *caenorhabditis elegans* | P42573 |
| Cel_casp1b/4b/5b | *caenorhabditis elegans* | Q9TZP5-3 |
| Cel_casp1c/4c/5c | *caenorhabditis elegans* | G5EBM1 |
| Dre_casp2 | *Danio rerio* | GCA_000002035.4 |
| Dre_casp3/7 | *Danio rerio* | GCA_000002035.4 |
| Dre_casp3 | *Danio rerio* | GCA_000002035.4 |
| Dre_casp6b | *Danio rerio* | GCA_000002035.4 |
| Dre_casp6c | *Danio rerio* | GCA_000002035.4 |
| Dre_casp6a | *Danio rerio* | GCA_000002035.4 |
| Dre_casp7 | *Danio rerio* | GCA_000002035.4 |
| Dre_casp10a | *Danio rerio* | GCA_000002035.4 |
| Dre_casp10b | *Danio rerio* | GCA_000002035.4 |
| Dre_casp9 | *Danio rerio* | GCA_000002035.4 |
| Hsa_casp1 | *Homo sapiens* | P29466 |
| Hsa_casp2 | *Homo sapiens* | P42575 |
| Hsa_casp3 | *Homo sapiens* | P42574 |
| Hsa_casp4 | *Homo sapiens* | P49662 |
| Hsa_casp5 | *Homo sapiens* | P51878 |
| Hsa_casp8 | *Homo sapiens* | O15519 |
| Hsa_casp14 | *Homo sapiens* | P31944 |
| Hsa_casp7 | *Homo sapiens* | P55210-3 |
| Hsa_casp9 | *Homo sapiens* | P55211 |
| Hsa_casp6 | *Homo sapiens* | P55212 |
| Hsa_casp10 | *Homo sapiens* | Q92851 |
| Mmu_casp1 | *Mus musculus* | P29452 |
| Mmu_casp2 | *Mus musculus* | P29594 |
| Mmu_casp3 | *Mus musculus* | P70677 |
| Mmu_casp6 | *Mus musculus* | O08738 |
| Mmu_casp7 | *Mus musculus* | P97864 |
| Mmu_casp10 | *Mus musculus* | O89110 |
| Mmu_casp9 | *Mus musculus* | Q8C3Q9 |
| Mmu_casp4b/5b | *Mus musculus* | P70343 |
| Mmu_casp4a/5a | *Mus musculus* | O08736 |
| Mmu_casp14 | *Mus musculus* | O89094 |
| Spu_casp3d/6d/7d | *Strongylocentrotus purpuratus* | GCA_000002235.4 |
| Spu_casp3e/6e/7e | *Strongylocentrotus purpuratus* | GCA_000002235.4 |
| Spu_casp1b/4b/5b | *Strongylocentrotus purpuratus* | GCA_000002235.4 |
| Spu_casp1c/4c/5c | *Strongylocentrotus purpuratus* | GCA_000002235.4 |
| Spu_casp3a/6a/7a | *Strongylocentrotus purpuratus* | GCA_000002235.4 |
| Spu_casp3b/6b/7b | *Strongylocentrotus purpuratus* | GCA_000002235.4 |
| Spu_casp6 | *Strongylocentrotus purpuratus* | GCA_000002235.4 |
| Spu_casp3/7 | *Strongylocentrotus purpuratus* | GCA_000002235.4 |
| Spu_casp1a/4a/5a | *Strongylocentrotus purpuratus* | GCA_000002235.4 |
| Spu_casp3c/6c/7c | *Strongylocentrotus purpuratus* | GCA_000002235.4 |
| Spu_casp10 | *Strongylocentrotus purpuratus* | GCA_000002235.4 |
| Sko_casp3a/7a | *Saccoglossus kowalevskii* | GCA_000003605.1 |
| Sko_casp3c/7c | *Saccoglossus kowalevskii* | GCA_000003605.1 |
| Sko_casp3d/7d | *Saccoglossus kowalevskii* | GCA_000003605.1 |
| Sko_casp3/6/7 | *Saccoglossus kowalevskii* | GCA_000003605.1 |
| Sko_casp10b | *Saccoglossus kowalevskii* | GCA_000003605.1 |
| Sko_casp6 | *Saccoglossus kowalevskii* | GCA_000003605.1 |
| Sko_casp3b/7b | *Saccoglossus kowalevskii* | GCA_000003605.1 |
| Sko_casp10a | *Saccoglossus kowalevskii* | GCA_000003605.1 |
| Sko_casp10c | *Saccoglossus kowalevskii* | GCA_000003605.1 |
| Bbe_casp3/6/7 | *Branchiostoma belcheri* | A0A6P5AKF3 |
| Bbe_casp3/7 | *Branchiostoma belcheri* | A0A6P4ZZL3 |
