## Supplementary File 3 for "Induction of apoptosis by double-stranded RNA was present in the last common ancestor of cnidarian and bilaterian animals"

**Supplementary file 3.** Detailed information on each of the studies used in cross-species comparative transcriptomic analysis

**Study 1**

The paper "A Sustained Immune Response Supports Long-Term Antiviral Immune Priming in the Pacific Oyster, Crassostrea gigas (Lafont *et al*., 2020)" focuses on the innate immune priming in oysters, particularly their response to Ostreid herpesvirus 1 (OsHV-1), a major pathogen. Its objective was to investigate the molecular basis of long-term resistance in Pacific oysters to Paciﬁc oyster mortality syndrome (POMS) following immune priming. Oysters were primed with poly(I·C), a viral mimic, and then exposed to OsHV-1. The study used transcriptomic analysis (RNA sequencing) to observe gene expression changes. Poly(I·C) induced a sustained upregulation of immune genes, notably those involved in interferon pathways and apoptosis. Our in-depth analysis into the differential expression of genes associated with the apoptosis pathway and their ontology analysis was sourced from Supplementary Files 3, 4, and 5. These files are instrumental in offering a comprehensive overview and detailed information critical to our comparative analysis (Figure 5).

**Study 2**

The paper "Induced Immune Reaction in the Acorn Worm, *Saccoglossus kowalevskii*, Informs the Evolution of Antiviral Immunity (Tassia et al., 2023)" presents a detailed study on the immune response of the acorn worm *Saccoglossus kowalevskii* to synthetic viral double-stranded RNA analog poly(I:C), providing insights into the evolution of antiviral immunity in Deuterostomia. The researchers injected *S. kowalevskii* with poly(I:C) to simulate viral infection, analyzing gene expression across different body regions at multiple time points post-injection. The study identified 423 genes that were differentially expressed in response to poly(I:C). The research showed the worm's immune system could recognize and react to the poly(I:C), indicating a complex innate immune response. The study used various bioinformatics tools for functional annotation and enrichment analysis. This revealed dynamic changes in immune response regulation, including genes associated with canonical innate immunity signaling pathways (like nuclear factor κB and interferon regulatory factor signaling) and metabolic processes (like lipid metabolism). We utilized the Supplementary Data, accessible through Zenodo at DOI: 10.5281/zenodo.7758058, to meticulously identify and analyze the expression patterns of genes linked with the apoptosis pathway. Additionally, this data aided in pinpointing the biological process Gene Ontology (GO) terms that are predominantly overrepresented among the up-regulated genes at each evaluated time point.

**Study 3**

The paper titled "Functional Analysis of ADARs in Planarians (Bar Yaacov, 2022)" explores the role of ADAR (Adenosine Deaminases Acting on RNA) proteins in planarian flatworms and aims to understand their evolutionary function, particularly in suppressing double-stranded RNA (dsRNA) responses. The researchers identified two ADAR homologs, ADAR1 and ADAR2, in the planarian species *Schmidtea mediterranea*. These homologs are evolutionarily distant from those in traditional lab models like flies and nematodes. The study used RNA interference (RNAi) to knock down ADAR1 and ADAR2 in planarians, observing the resultant effects. Knockdown of ADAR1, but not ADAR2, caused lesions and lethality in planarians, indicating its essential role. RNA sequencing (RNA-Seq) analyses revealed significant changes in gene expression upon knockdown of ADAR1 and ADAR2. Notably, genes involved in dsRNA responses were up-regulated. Knockdown of ADAR1 and ADAR2 led to a reduction in the number of cells infected with a dsRNA virus, suggesting that ADARs suppress a bona fide anti-viral dsRNA response. This paper provides significant insights into the evolutionary role of ADARs in planarians and their importance in regulating immune responses to dsRNA, which can have implications for understanding viral defense mechanisms across different species. We utilized S2 Table RNA-Seq differential expression analysis and S3 Table KEGG pathway analysis of this research to examine whether apoptotic pathway genes were upregulated and to assess the significance of apoptosis-related terms.

**Study 4**

The paper titled "Exploring gene expression changes in the amphioxus gill after poly(I:C) challenge using digital expression profiling (Zhang et al., 2017)" investigates the response of amphioxus gills to immune challenges, particularly focusing on gene expression changes following exposure to poly(I:C), a viral mimic. Amphioxus is a basal chordate and serves as a model organism for understanding vertebrate evolution, especially in terms of immune system development. Gene expression levels were quantified, and differentially expressed genes (DEGs) were identified. Gene Ontology (GO) and Kyoto Encyclopedia of Genes and Genomes (KEGG) enrichment analyses were conducted on these DEGs. The study identified several key genes responding to the immune challenge, indicating that the amphioxus gill participates in antiviral immunity. Due in this study the supplementary files were not well detailed, we reanalyzed the data to identify the upregulated genes and the significant terms across various treatments. The data underpinning the findings of this study are available in the NCBI Sequence Read Archive (SRA). The relevant datasets can be accessed using the following run accession numbers: SRR5980103, SRR5980104, SRR5980106, SRR5980107, SRR5980108, and SRR5980109.

**Study 5**

The paper titled "A minimal RNA ligand for potent RIG-I activation in living mice" by Melissa M. Linehan and colleagues (2018) focuses on the development of potent synthetic activators of the vertebrate immune system, specifically targeting the RIG-I receptor. These activators are short triphosphorylated stem-loop RNAs (SLRs), which, when introduced into mice, induce a potent interferon response and activate genes essential for antiviral defense. RNA sequencing was conducted to analyze gene expression in mouse spleen following SLR and poly(I:C) injection. This provided insights into the distinct gene expression patterns induced by each. SLRs predominantly induced type I interferon genes, while poly(I:C) induced a different set, including type III interferons. We analyzed tables S1 and S2 to identify genes associated with apoptosis and used these gene lists to investigate significant GO terms using **Go**rilla (Eden et al., 2009-https://pubmed.ncbi.nlm.nih.gov/19192299/).

**Study 6**

The paper "Gene expression variability across cells and species shapes innate immunity (Hagai *et al*., 2018)" provides a comprehensive analysis of how the innate immune response varies across different cells and species, focusing on the transcriptional divergence and variability in gene expression. The study focuses on the innate immune response, the first line of defense against pathogens. It is characterized by a broad range of cell-to-cell variability and transcriptional divergence across species. The innate immune response includes upregulation of antiviral and inflammatory cytokines, gene upregulation to restrict pathogens, and induction of cell death. The research employed bulk and single-cell transcriptomics in fibroblasts and mononuclear phagocytes from various species, stimulated with immune stimuli. The study's primary objective was to map the architecture of the innate immune response and understand its evolution across different species and cellular variability. There was a notable divergence in the transcriptional response of fibroblasts to dsRNA (poly(I:C)) across species (humans, macaques, mice, and rats). The divergence was also observed in mononuclear phagocytes stimulated with LPS (lipopolysaccharide). This divergence was quantified by assessing the fold-change in gene expression and its relation to the phylogenetic relationship between species. Using only data from fibroblast cells (Supplementary Tables 3 and 4), we determined whether up-regulated genes were linked to the apoptosis pathway and employed these lists of differentially expressed genes to identify GO terms related to apoptosis in human and mouse.

**Study 7**

The paper titled "Functional Characterization of the Cnidarian Antiviral Immune Response Reveals Ancestral Complexity (Lewandowska et al., 2021)" investigates the antiviral immune response of the cnidarian *Nematostella vectensis*, particularly focusing on the function of the retinoic acid-inducible gene I (RIG-I)-like receptors (RLRs). These receptors are known to detect viral double-stranded RNA (dsRNA) in bilaterians but activate different antiviral pathways in vertebrates and nematodes. The study demonstrates that polyinosinic:polycytidylic acid (poly(I:C)), a mimic of long viral dsRNA and a primary ligand for the vertebrate RLR melanoma differentiation-associated protein 5 (MDA5), triggers a complex antiviral immune response in *N. vectensis*, showing characteristics of both vertebrate and invertebrate systems. We reanalyzed the raw data from this study, with our methodology detailed in the main text.
